# From heterogenous morphogenetic fields to homogeneous regions as a step towards understanding complex tissue dynamics

**DOI:** 10.1101/696252

**Authors:** Satoshi Yamashita, Boris Guirao, François Graner

## Abstract

Within developing tissues, cell proliferation, cell motility, and other cell behaviors vary spatially, and this variability gives a complexity to the morphogenesis. Recently, novel formalisms have been developed to quantify tissue deformation and underlying cellular processes. A major challenge for the study of morphogenesis now is to objectively define tissue sub-regions exhibiting different dynamics. Here we propose a method to automatically divide a tissue into regions where the local deformation rate is homogeneous. This was achieved by several steps including image segmentation, clustering, and region boundary smoothing. We illustrate the use of the pipeline using a large dataset obtained during the metamorphosis of the *Drosophila* pupal notum. We also adapt it to determine regions where the time evolution of the local deformation rate is homogeneous. Finally, we generalize its use to find homogeneous regions for the cellular processes such as cell division, cell rearrangement, or cell size and shape changes. We also illustrate it on wing blade morphogenesis. This pipeline will contribute substantially to the analysis of complex tissue shaping and the biochemical and bio-mechanical regulations driving tissue morphogenesis.

**Summary statement:** Tissue morphogenesis is driven by multiple mechanisms. This study proposes a methodology to identify regions in the developing tissue, where each of the regions has distinctive cellular dynamics and deformation.

## 2 Introduction

During tissue development, morphogenesis is accompanied by cellular processes such as cell division, cell rearrangement, cell size and shape changes, apical constriction, and apoptosis. The cellular processes are coordinated together, yielding collective cell migration and local deformation of each tissue region, resulting in convergent extension or epithelial folding. Furthermore, the local deformations of different tissue regions are coordinated too, resulting in large scale tissue morphogenesis. Coordinations between invaginated mesoderm and covering ectoderm (Rauzi et al., 2015; Perez-Mockus et al., 2017), between invaginated midgut and elongated germ-band (Collinet et al., 2015; Lye et al., 2015; Dicko et al., 2017) of *Drosophila* embryo, between contracting wing hinge and expanding wing blade in *Drosophila* pupa (Etournay et al., 2015; Ray et al., 2015), or between invaginated neural plate and covering epidermal ectoderm of *Xenopus* embryo (Brodland et al., 2010), provide examples of how mechanical force generated in one region can drive large scale deformation in adjacent regions. In these cases, the regions which behave differently are easily distinguished by specific gene expressions.

However, many tissues were found to be heterogeneous but without obvious boundary between such regions, leaving analysis limited to arbitrary regions drawn as a grid parallel to tissue axes, or regions expressing already known differentiation maker genes. Measured tissue deformation rate showed a large heterogeneity (accompanied by a heterogeneity in cellular processes such as cell proliferation rate, cell division, cell rearrangement, change of cell shape), and smooth spatial variations across the tissue, in *Drosophila* notum in a developing pupa (Bosveld et al., 2012; Guirao et al., 2015) (Fig. 1A, B), *Drosophila* wing blade (Etournay et al., 2015), blastopore lip of *Xenopus* gastrula (Feroze et al., 2015), chick gastrula (Rozbicki et al., 2015; Firmino et al., 2016), mouse palatal epithelia (Economou et al., 2013), and mouse limb bud ectoderm (Lau et al., 2015). Recent formalisms have enabled us to measure and link quantitatively cellular processes with tissue deformation (Blanchard et al., 2009; Guirao et al., 2015; Etournay et al., 2015; Merkel et al., 2017). Those studies showed that cellular quantities also vary smoothly across the tissue. In addition, the causal relationship between cellular processes and tissue deformation is not always trivial, making it difficult to identify regions those actively drive morphogenesis and that passively deformed by adjacent regions.

**Fig. 1:**
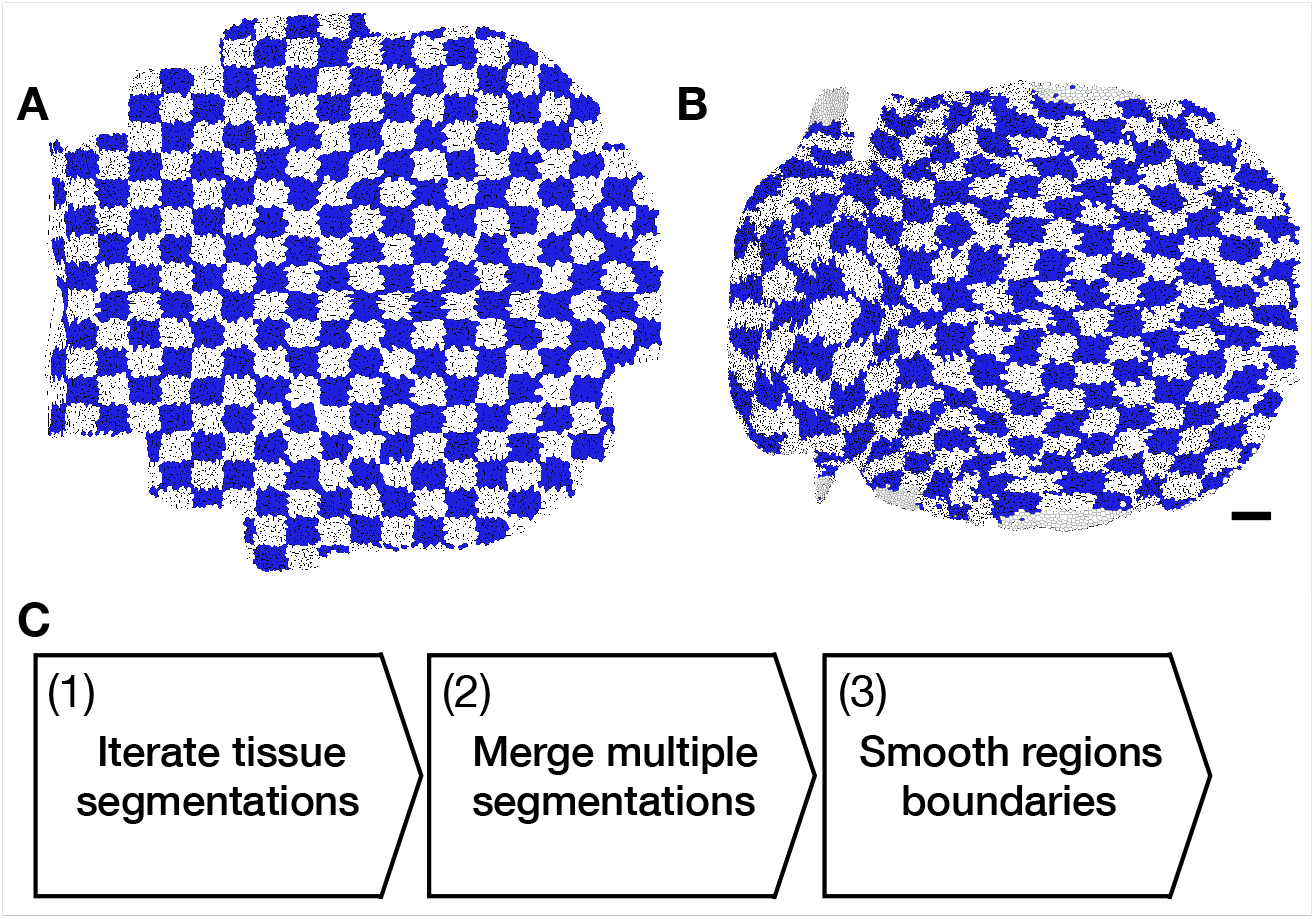
Morphogenesis of *Drosophila* pupa notum and overview of tissue segmentation pipeline. (**A**, **B**) Heterogeneity of tissue morphogenesis. Here a *Drosophila* notum at 12 hr after pupa formation (APF), with arbitrary regions drawn from a grid (**A**), and at 32 hr APF, showing the heterogeneous deformation of previous regions using cell tracking (**B**). Cell patches are shown with blue and white check pattern. (**C**) Pipeline of the tissue segmentation. (1) Iteration of fast tissue segmentation with random seeding, using region growing algorithm. (2) Merging multiple tissue segmentations of step 1 into a single objective tissue segmentation, using label propagation algorithm on a consensus matrix. (3) Smoothing regions boundaries resulting of step 2, using cellular Potts model.

To study the spatial regulation of morphogenesis at tissue scale, we developed a new multi-technique pipeline to divide a tissue into sub-regions based on quantitative measurements of static or dynamic properties of cells or tissues. Our tissue segmentation pipeline consists of two steps and an optional third step: a first fast tissue segmentation attempted several times with random seeding, then merging these multiple tissue segmentations into a single one, then if necessary smoothing the resulting regions boundaries (Fig. 1C). In contrast to other image segmenting methods like the watershed algorithm, this method is designed to accept any kind of quantity of biological interest, not only a scalar but also a vector, a tensor, and combination of them. We apply it to the morphogenesis of *Drosophila* pupa dorsal thorax and wing blade. They were divided based on the tissue deformation rate or how the cellular processes contribute to the tissue deformation. Obtained sub-regions showed distinctive patterns of deformation and cellular processes with higher homogeneity than those along tissue axes. Interestingly, the tissue segmentations based on the local tissue deformation rate and on the cellular processes included some similar regions, suggesting that the cellular processes were regulated similarly inside the regions, therefore resulting in homogeneous tissue deformations inside those regions.

## 3 Results I : Development of automatic tissue segmentation algorithm

### 3.1 Image segmentation by region growing algorithm

Finding distinctive and homogeneous regions inside the heterogeneous tissue amounts to segmenting the geometrical space while keeping the points inside each region as similar as possible to each other in the property space. Here, we call *property space* any morphogenesis quantification measured in the tissue, whereas *geometrical space* refers to the two-dimensional space of cell patch positions inside the tissue.

Given a set of objects, collecting similar objects to divide them into groups is generally a task of cluster analysis. However, the cell patches distribute both in the property space and geometrical space. On the assumption that expression patterns of genes responsible for morphogenesis make connected regions, and to study physical interactions between the regions, we aimed at getting connected regions. The initial tissue segmentation first defines a metric of similarity between cells, and then a tissue is divided into regions containing similar cells. The image segmentation tool, called region growing (Adams and Bischof, 1994; Ma et al., 2010) (Fig. 2A), was inspired by a study segmenting mouse heart based on cell polarity (Le Garrec et al., 2013).

**Fig. 2:**
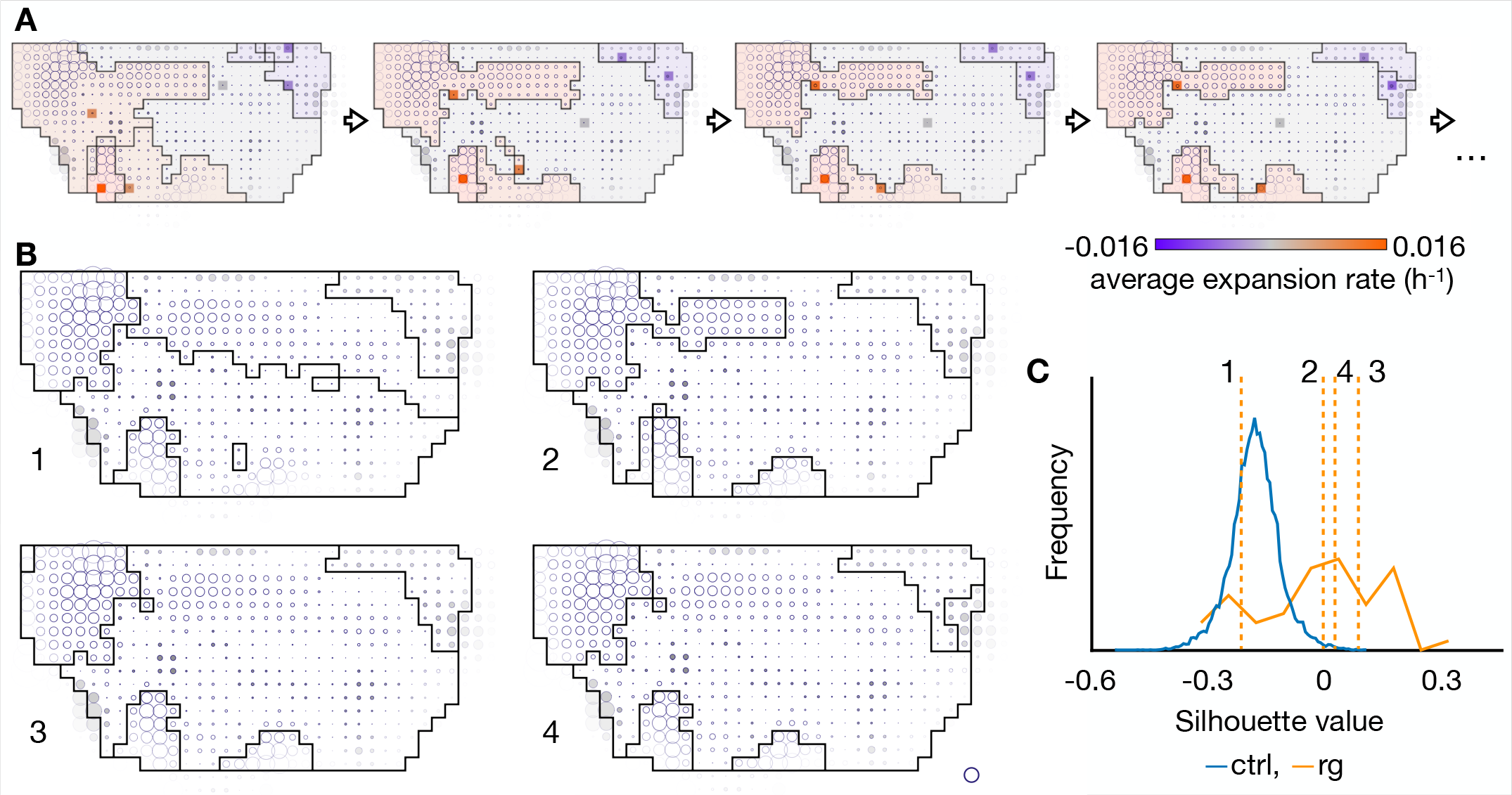
Tissue segmentation by region growing algorithm. Cell patches expansion/contraction rates are represented by size of white/gray circles. (**A**) Process of region growing algorithm. Points of given number (6 in the shown example) are chosen randomly as initial seeds of regions, and the regions are expanded by collecting points similar to the seeds from their neighbors. Once the field is segmented, the seeds are updated to region centroids in the geometrical space and means in the property space, and the expansion of the regions are performed again from the new seeds. The seeds are shown with colored square, where the color represents an expansion rate of the regions. The regions are colored lighter for visibility. This update of the seeds and the regions are iterated until it reaches a convergence. (**B**) Four example results of region growing. (**C**) Histogram of silhouette value: blue for control segmentations, orange for region growing. Dotted vertical orange lines show silhouette values of the four examples shown in **B**. For clarity, in this figure and others, frequency axis units are not included.

To validate the algorithm, we first tested segmentation on a simple example, namely the change in cell patch areas from 12 to 32 hr APF (Fig. 2B). The overall change in cell patch areas defines the total tissue growth, while spatially heterogeneous changes in cell patch areas result in local deformation, changes in tissue region proportions, and overall tissue shape change. Technically speaking, the change in cell patch areas is a scalar field, defined as the trace of the tissue deformation rate tensor. The region growing succeeded in finding expanding regions in posterior, lateral posterior, and lateral parts and a shrinking region in anterior part.

However, the results varied dependent on the initial seeds. In contrast to a segmentation of immuno-stained image, where a true segmentation is well defined, the morphogenetic properties vary continuously with space, making it difficult to determine and validate the resultant segmentations. The silhouette, a measurement of region homogeneity (the silhouette of an object would be 1 if it was similar to all objects in the same cluster, and *−*1 if it was more similar to objects in other clusters), differed from one segmentation to the other (Fig. 2C). To assess the significance of the homogeneity, we compared it with the average silhouette of randomly made control segmentations. Some of the region growing results had a low silhouette, even lower than that of half of the control segmentations (Fig. 2C), which means they were lacking any signification.

Among the various results because of the random initial seeding, we don’t know which one should be compared with gene expression patterns or fed forward to a study of mechanical interactions between the regions. For practical applications, we need a single segmentation result for a given morphogenetic property.

### 3.2 Defining a single tissue segmentation using label propagation on a consensus matrix

To obtain a single tissue segmentation, we turned to consensus clusterings. In fact, since resultant segmentations of the region growing were dependent on randomly given initial values, we ran multiple trials and merged multiple segmentation results into a single one. Given multiple partitions, the consensus clustering returns the partition which is the most similar to all of the initial partitions. We tried several consensus clustering algorithms, and found the *label propagation on a consensus matrix* (Lancichinetti and Fortunato, 2012; Raghavan et al., 2007) returning regions similar to the results of region growing.

The label propagation on a consensus matrix converted multiple tissue segmentations into a weighted graph where weight of an edge represented a frequency of segmentations in which incident vertices (points) belonged to the same region (Fig. 3A). Then labels on the vertices were propagated according to the weight so that the same label was assigned to points which were frequently included in the same region among the given multiple region growing segmentations.

**Fig. 3:**
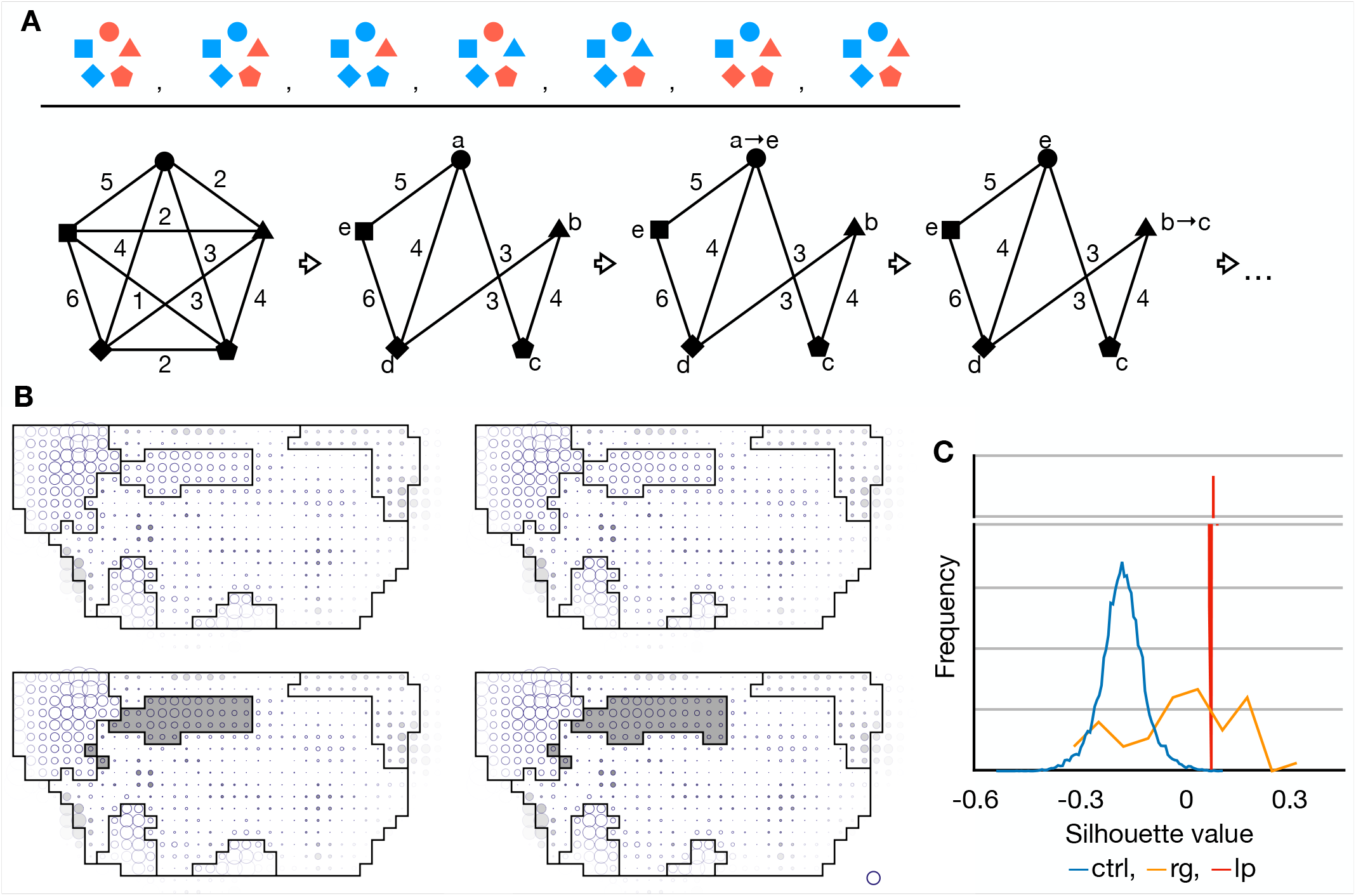
Tissue segmentation by label propagation on a consensus matrix. (**A**) Process of label propagation algorithm. Multiple clusterings (upper) are converted to a consensus matrix, which gives weights to a complete graph on the objects being clustered (lower left). Edges with weights less than a given threshold are removed. All objects are initially assigned labels different to each other. And then, one by one in random order, each label is updated to the most frequent one weighted by edges incident to the object until it reaches a convergence. (**B**) Four example results of label propagation on the same consensus matrix. (**C**) Histogram of silhouette value: blue for control segmentations, orange for region growing, red for label propagation.

The label propagation returned results similar to the region growing segmentations (Fig. 2B, 3B). Also, the label propagation results were more similar to each other than results of region growing, assessed with adjusted Rand indices (ARI), a measurement of similarity between two partitions (ARI of identical partitions would be 1). ARI were 0.50 *±* 0.21 among the results of the region growing and 0.97 *±* 0.02 among the results of the label propagation. They showed similar average silhouette values, similar to median of those of region growing results, but smaller than the highest value of those of region growing (Fig. 3C). The average silhouette of the label propagation result was higher than those of 99.95% of the randomly made control segmentations.

However, a consensus clustering algorithm ignores original properties of objects in principle and divides the objects only based on how they were divided among given partitions, and thus it might return disconnected regions and zigzag boundary between them. Some segmentations in Figure 3B also included disconnected regions as marked by gray color.

### 3.3 Smoothing of tissue segmentation results by cellular Potts model

For the case of complex boundary and disconnected points, we prepared an optional step to smooth the boundary and remove disconnected points when needed. To smooth the consensus regions boundaries, we employed cellular Potts model, which simulates dynamics of a cellular tissue by calculating energies of cells from their geometry, trying to decrease the total energy. In our application to the boundary smoothing, the energy was lower when the region boundary was shorter and the homogeneity was higher (Fig. 4A). The boundary length and homogeneity were balanced so that all regions had enough smooth boundary, evaluated by a circularity (Bosveld et al., 2016), and was as homogeneous as possible. The result was not affected largely by extending a duration of the simulation, and thus we stopped the simulation when it was enough smoothed.

**Fig. 4:**
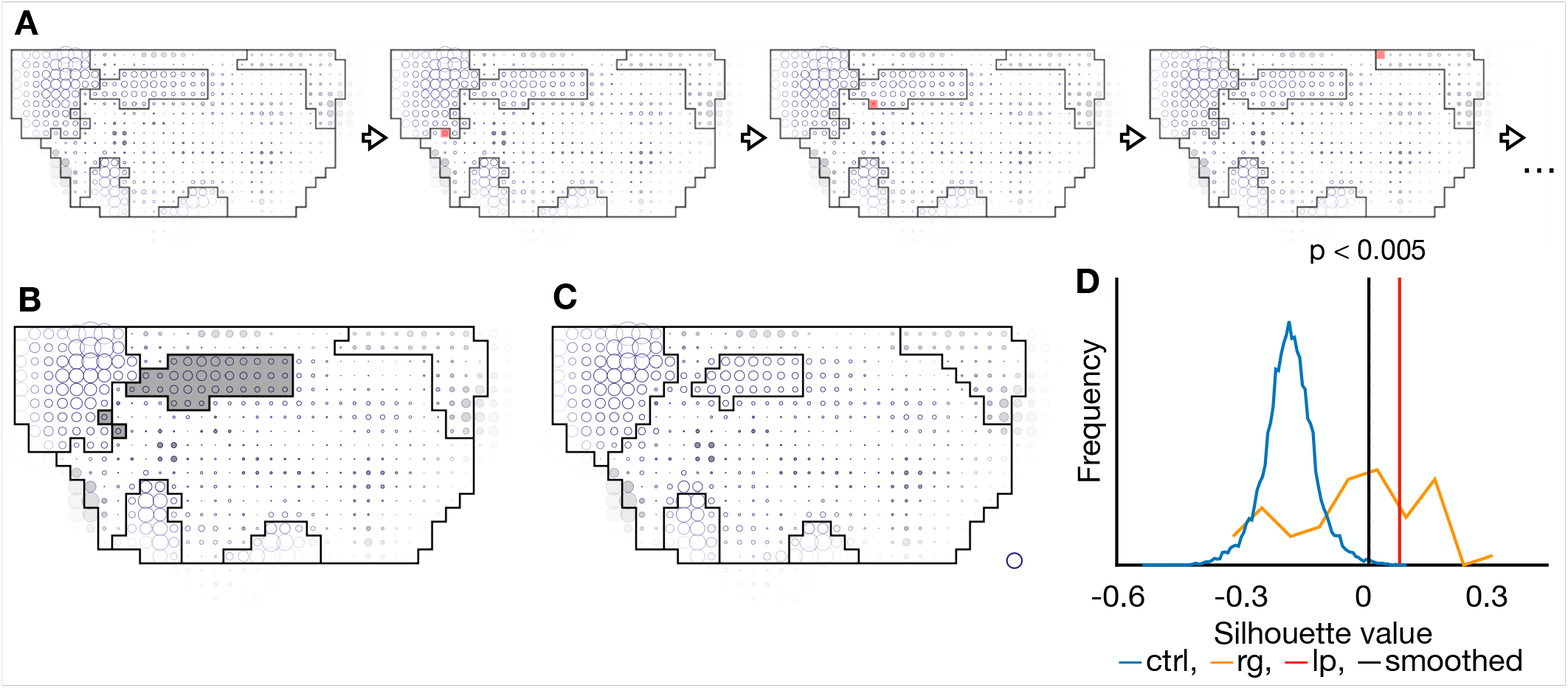
Boundary smoothing by cellular Potts model. (**A**) Process of cellular Potts model. A pixel is randomly chosen and changes its belonging region if it decreases boundary length and/or increases homogeneity (marked by red). (**B**) Result of label propagation with a disconnected region shown by gray color. (**C**) Result of boundary smoothing by cellular Potts model. (**D**) Histogram of silhouette value: blue for control segmentations, orange for region growing, red vertical line for label propagation, and black vertical line for regions smoothed by cellular Potts model. Dotted blue line shows threshold for the highest 0.5% of the control segmentations.

It smoothed boundaries and removed disconnected cell patches (Fig. 4B, C) while keeping the average silhouette value higher than those of 99.5% of the randomly made control segmentations (Fig. 4D). Since the cellular Potts model implementation includes the Metropolis update, i.e., choosing a pixel randomly and updating the pixel by probability according to a change of the energy, resultant smoothed segmentations varied among different trials even with the same parameters and initial segmentation. Therefore we iterated the cellular Potts model smoothing 50 times and integrated its results by the label propagation algorithm again.

Now we have a pipeline of the region growing, the label propagation, and the optional cellular Potts model to divide a field of property (scalar, tensor, or any kind of value with metric) into regions. The resultant regions are homogeneous, where points in each region are more similar to each other than to points in other regions.

## 4 Verification of the method against simulated data

We tested our method whether it can segment a simulated tissue using a normal cellular Potts model.

We prepared a 2D tissue model in which two types of cells with different surface tensions were assembled (Fig. 5A). The tissue was geometrically compressed in the horizontal direction and extended in the vertical direction, so that the cells retained their area but they are vertically elongated (Fig. 5B). Because of the surface tension, the cells tried to minimize their perimeter, and thus they were rearranged and rounded (Fig. 5C, Movie 1). The tissue was split in a 4 *×* 7 grid, and the deformation and cellular processes after the compression were measured in each cell patch. The measured deformation and cellular processes were averaged among 12 simulations. The cells were rearranged similarly between the two types of cells (Fig. 5D, Movie 1). However, when we tried the segmentation based on the cell rearrangement, the cells were successfully distinguished (Fig. 5E). When the tissue was segmented into three regions, it included a small and disconnected region after the first label propagation and smoothing (Fig. 5F, G). The cell rearrangement rate was slightly higher among the cells with lower surface tension (Fig. 5H). In general, lower surface tension would allow larger fluctuation, and the larger fluctuation might have facilitated the faster cell rearrangement. The obtained two regions showed a significantly high homogeneity (Fig. 5I). With these results, we confirmed that our method could properly identify groups of cells with different mechanical properties based on their behavior.

**Fig. 5:**
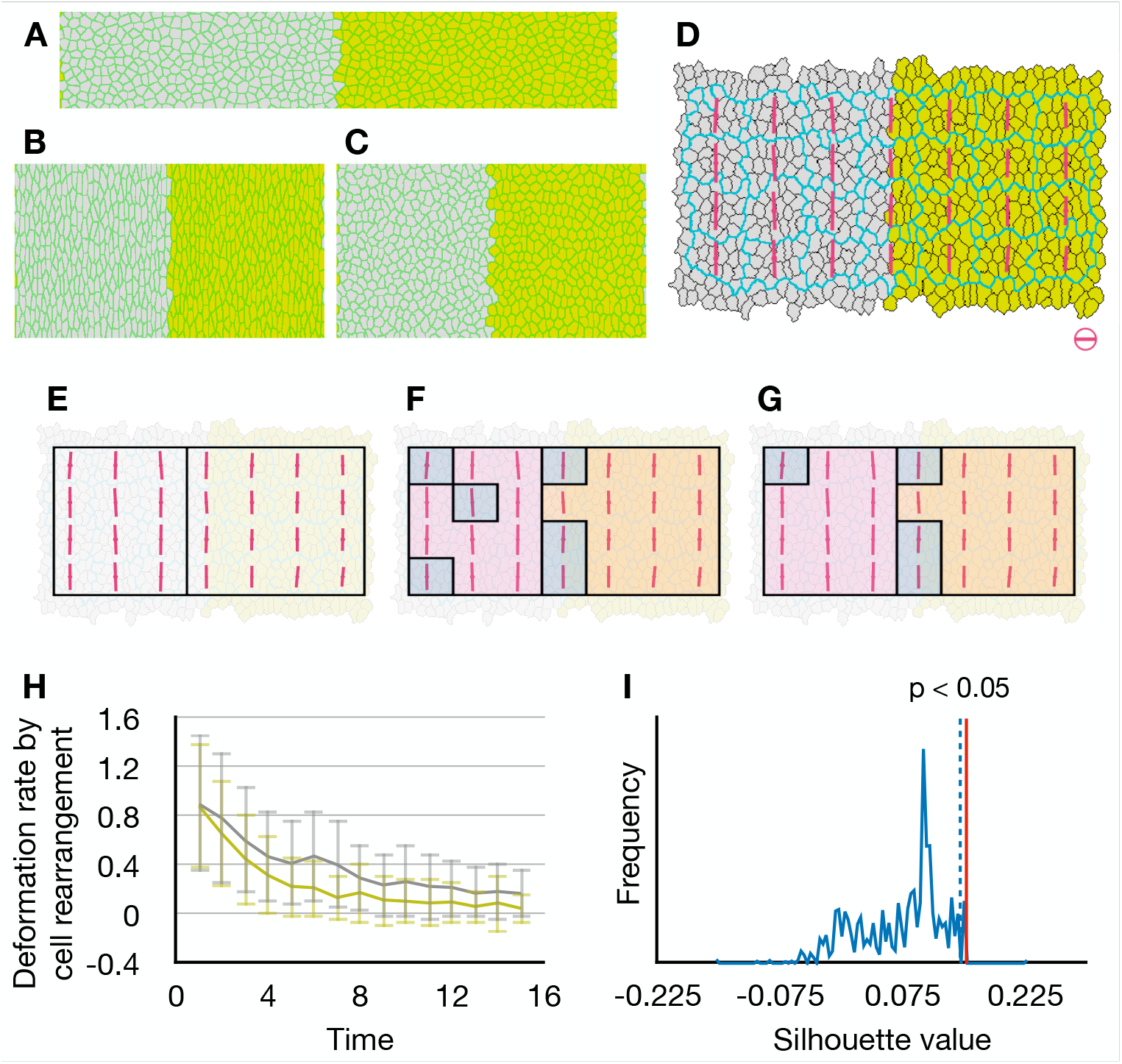
Segmentation of simulated tissue. (bf A) Initial configuration of the model tissue. The cells with lower surface tension were colored gray, and the cells with higher surface tension were colored yellow. Cell perimeters were colored green. (**B**, **C)** Tissue after the compression (**B**) and after the simulation (**C**). (**D**) Cell rearrangement in each cell patch after the compression. Pale blue line shows the cell patches. Magenta bars represent direction and rate of assumptive tissue deformation caused by the cell rearrangements. Scale bar and circle indicate deformation rate 0.2 h*^−^*^1^. (**E**) Segmentation into two regions based on the cell rearrangements. It is a result of the first label propagation, but it did not need smoothing. (**F**, **G**) Segmentation into three regions based on the cell rearrangements, after the first label propagation (**F**) and the boundary smoothing (**G**). The regions were colored for visibility. (**H**) Plot of the assumptive deformation rate caused by the cell rearrangements. Eigenvalue of the tensor for the cell rearrangements represents a rate of tissue elongation by the cell rearrangement, and it is plotted against time after the compression. Gray line and yellow line show the average rate among the cells with lower surface tension and higher tension respectively. Error bars show their standard deviation. (**I**) Histogram of average silhouette value of control segmentations divided into two regions. Red vertical line shows silhouette value of label propagation result (**E**). Dotted blue line shows threshold for the highest 5% of the control segmentations.

## 5 Results II : Tissue segmentation based on tissue morphogenesis

We now turn to property spaces better representing tissue morphogenesis. In Guirao et al. (2015), tissue deformation rate (**G**) and underlying cellular processes, cell division (**D**), cell rearrangement (**R**), cell shape change (**S**), and cell delamination (**A**) were quantified into tensors. The tensors were obtained from change of the texture averaged over 20 hr from 12 hr APF to 32 hr APF or over 2 hr at each time point. By comparing the tensors, for example, one can check whether cell divisions and cell rearrangements elongated tissue in the same direction or attenuated each other. In the same way, by comparing the tensors of deformation rates with a unit tensor which has the same direction of elongation as tissue deformation rate, one can estimate an amplitude of the tissue deformation rate and how much the cellular processes contribute to the tissue deformation in both terms of contraction/expansion (isotropic deformation) and narrowing/elongation (anisotropic deformation) (Guirao et al., 2015). They are scalar value and denoted by **G***_//_* for the tissue deformation rate, **D***_//_* for cell division, **R***_//_* for cell rearrangement, **S***_//_* for cell shape change, and **A***_//_* for cell delamination. For the sake of clarity, we call the tissue deformation rate and the cellular processes averaged over the whole 20 hr from 12 to 32 hr APF *time-average* tissue deformation rate and cellular processes.

The effective contributions averaged over the whole tissue showed dynamic time evolution (Fig. S1), with a large peak of cell division and cell shape change around 16 hr APF, second small wave of cell division around 22 hr APF, and gradual increase of cell shape change and cell rearrangement. The effective contributions also showed large variance across the tissue at each time point. Therefore we included the time evolution in the property space. Assume that there are two regions in a tissue where the tissue expands, the first region expands during 14-17 hr APF, and the second region expands during 25-28 hr APF, resulting in similar size changes, then the two regions cannot be distinguished by the time-average expansion rate. To distinguish them, we compared a property at each time point and summed up its difference through the whole time. When two cell patches always behaved similarly, then the difference at each time point is small and so the total difference is small too, whereas cell patches with deformations occurring in different timing are separated at each time point and thus the total difference gets large. In contrast with time-average, we call the sum of difference at each time point *time-evolution*.

### 5.1 Tissue segmentations based on tissue deformation rate and cellular processes effective contributions

We first divided the tissue based on time-average and time-evolution of tissue deformation rate. The similarity was given by Euclidean distance of tensors. The notum was divided into anterior-middle-posterior and medial-lateral regions by both the time-average and time-evolution, while the middle regions were smaller and the middle lateral region extended medially in the segmentation based on time-evolution (Fig. 6A-D).

**Fig. 6:**
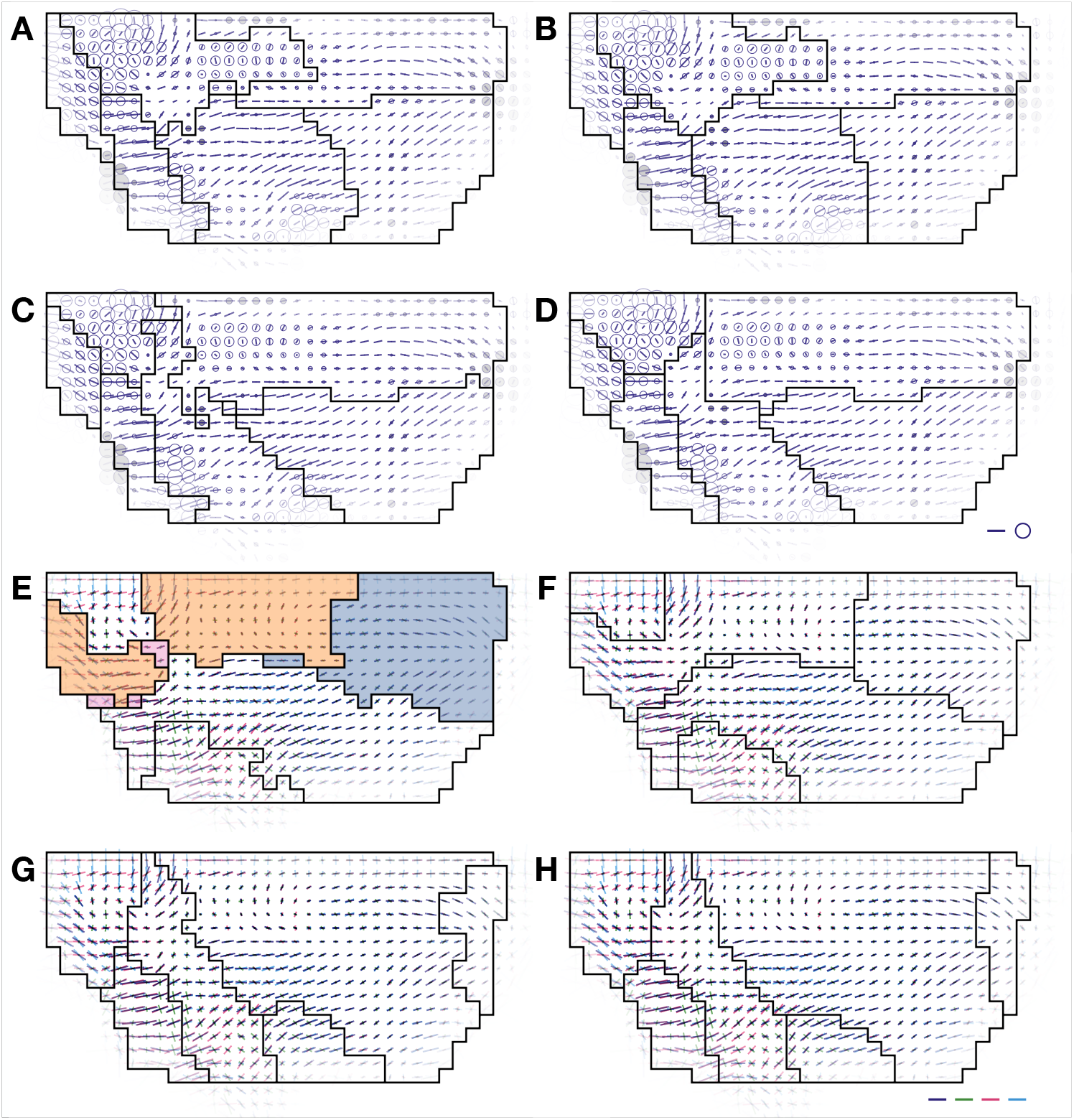
Segmentations based on tissue deformation and underlying cellular processes. For each cell patch, direction of elongation is represented by a bar, and the effective contributions of cellular processes are indicated by relative directions of deformation rate between the tissue and each cellular process. For quantification and representation of tissue deformation rate and cellular processes, see Methods and Guirao et al. (2015). (**A**-**H**) Segmentations based on time-average tissue deformation rate (**A**, **B**), time-evolution of deformation rate (**C**, **D**), time-average cellular processes effective contributions (**E**, **F**), and time-evolution of cellular processes (**G**, **H**). First column shows results of the first label propagation (**A**, **C**, **E**, **G**) and second column shows results of boundary smoothing (**B**, **D**, **F**, **H**).

Next, we divided the tissue based on time-average and time-evolution of cellular processes. The amplitude of tissue deformation rate and cellular processes effective contributions were combined in a vector, and their similarity was given by Euclidean distance of vector. In contrast to the segmentations based on the time-average and time-evolution of tissue deformation rate, the segmentations based on time-average and time-evolution of the cellular processes were dissimilar to each other (Fig. 6E-H). The segmentation based on time-evolution of cellular processes included a posterior region, a large anterior region, a neck-notum boundary region, lateral posterior region, a scutum-scutellum boundary region, and a lateral region (Fig. 6H).

While a change in the minimum circularity for the boundary smoothing did not affect the segmentation based on the time-average tissue deformation rate (Fig. S2A), shape of the boundary changed dependent on the minimum circularity for the segmentations based on the other three property spaces (Fig. S2B-D). The cellular Potts model returns regions with circularity higher than the given minimum value if it is the most homogeneous segmentation.

We also tried dividing the tissue into various numbers of regions (Fig. S3). In many cases, an increase of the number of the regions resulted in subdividing the regions already obtained. In addition, when the number was too large, some results of the first label propagation included small regions similar to one observed in the segmentation of simulated data (Fig. 5F). Those small regions were absorbed into surrounding regions during the smoothing by cellular Potts model. Then the final label propagation tried to integrate regions smaller than the final segmentation, returning small and sometimes disconnected regions (Fig. S3 third column below third row and fourth column below sixth row), again like one of simulated data (Fig. 5G). Thus the existence of small regions suggest that it is over-segmented.

### 5.2 Correspondence between segmentations based on cellular processes and tissue deformation rate

Both of the segmentations based on time-evolution of tissue deformation rate and cellular processes effective contributions included the large anterior region, the middle boundary region, the lateral posterior region, and the posterior region, although the anterior and posterior regions were divided into medial and lateral subregions in the segmentation based on the tissue deformation rate. Figure 7A-D show overlap between segmentation based on time-evolution of cellular processes (Fig. 6H) and the others (Fig. 6B, D, F) or a conventional large grid parallel to tissue axes. The middle lateral and posterior lateral regions in the segmentation based on time-evolution of tissue deformation rate and the middle scutum-scutellum boundary region and lateral posterior regions in the segmentation based on time-evolution of cellular processes overlapped each other (Fig. 7C).

**Fig. 7:**
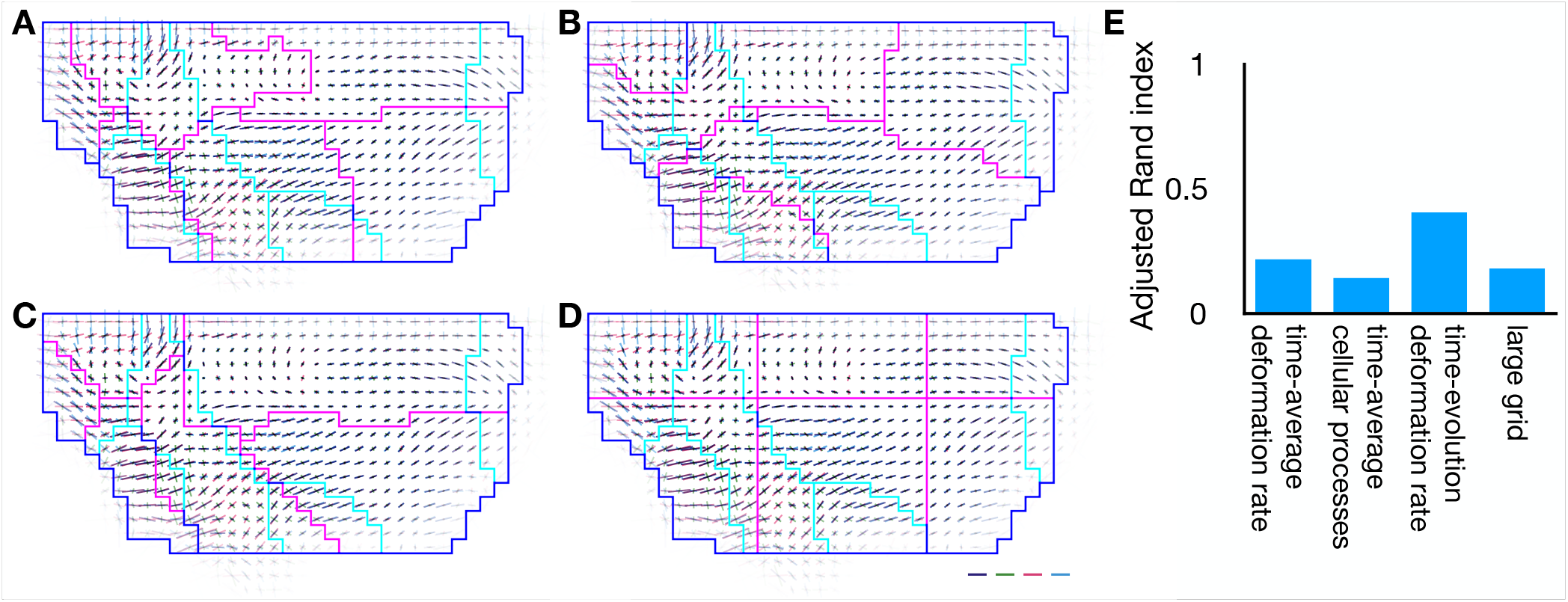
Correspondence between segmentations based on cellular processes and deformation rate. (**A**-**D**) Overlays of segmentations, where segmentation based on time-evolution of deformation rate is shown by cyan line, while segmentations based on time-average deformation rate (**A**), time-average cellular processes (**B**), time-evolution of cellular processes (**C**), and large grid (**D**) are shown by magenta line. (**E**) Adjusted Rand indices of **A**-**D**.

We also evaluated the overlap between the segmentations by ARI (Fig. 7E). Despite the difference between the anterior subregions, the segmentations based on time-evolution of tissue deformation rate and cellular processes overlapped each other more than the others.

### 5.3 Homogeneity of the obtained regions

Next, we evaluated the homogeneity of the obtained regions. The time-evolution of tissue deformation rate was similar among cells inside regions of the segmentations based on time-average and time-evolution of tissue deformation rate except the middle-lateral region of the former (Fig. 8A, B). On the other hand, the large grid segmentation showed large heterogeneity in the posterior regions (Fig: 8C). The average silhouette value of the segmentation based on the time-evolution of deformation rate was higher than those of 99.5% of the control segmentations (average silhouette for label propagation: 0.0568, for smoothed regions: 0.0600, the maximum of the smallest 99.5% of controls average silhouettes: 0.0425) (Fig. 8D). The average silhouette of the segmentation based on time-averaged tissue deformation rate was also higher than 95% of the control segmentations (average silhouette for label propagation: 0.0336, for smoothed regions: 0.0260, the maximum of the smallest 95% of controls average silhouettes: -0.0098). On the other hand, that of the conventional grid segmentation was close to median of the control segmentations (average silhouette: -0.0694).

**Fig. 8:**
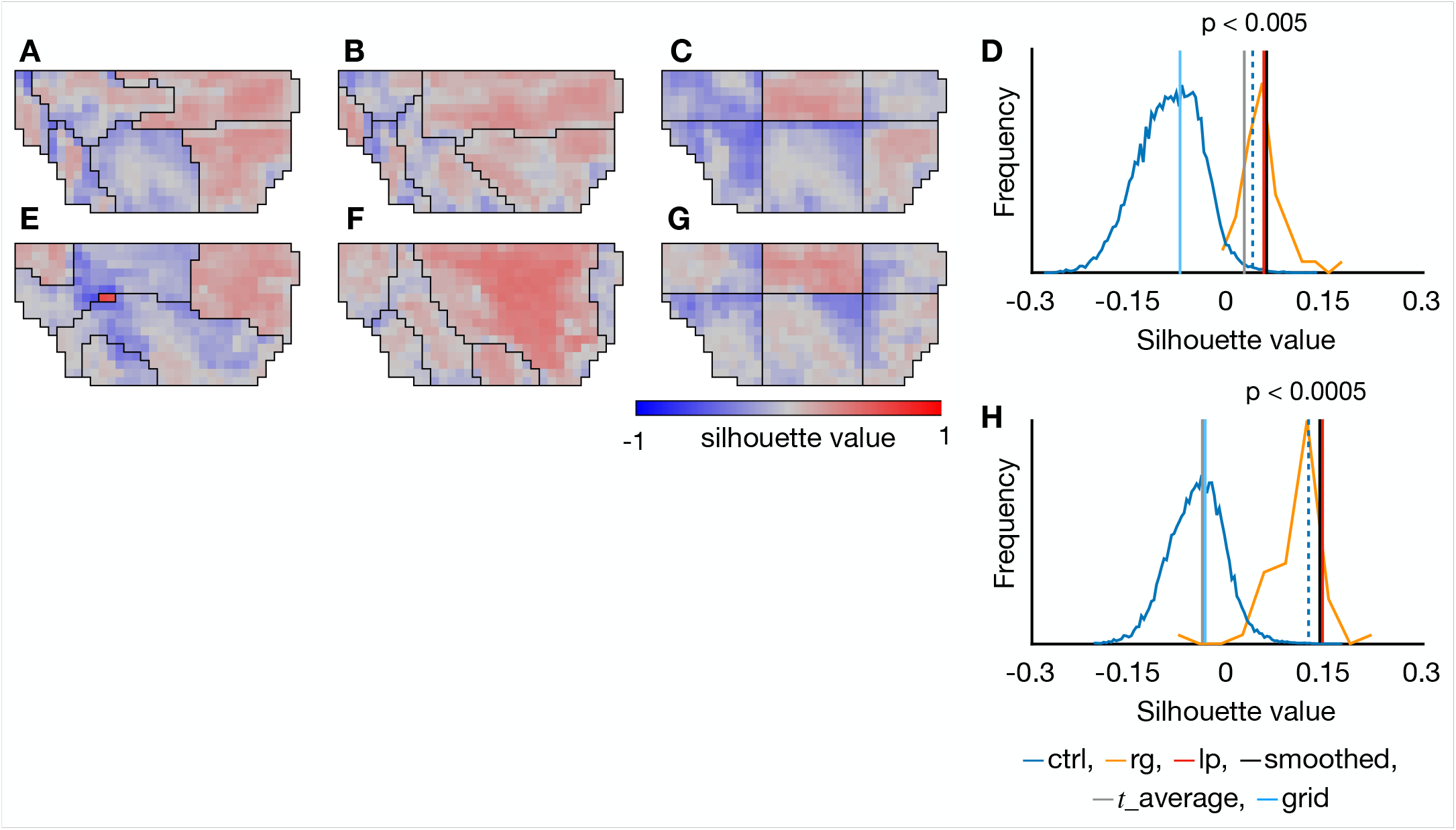
Homogeneity in the obtained regions. (**A**-**C**) Heat map of silhouette value measured with time-evolution of tissue deformation rate in segmentations based on time-average deformation rate (**A**), time-evolution of deformation rate (**B**), and large grid (**C**). (**D**) Histogram of silhouette value: blue for control segmentations, orange for region growing. Red vertical lines show silhouette value of label propagation results. Black vertical lines show silhouette value of regions smoothed by cellular Potts model. Dotted blue lines show threshold for the highest 0.5% of the control segmentations. Gray and cyan vertical line shows silhouette value of segmentation based on time-averaged deformation rate and large grid. (**E**-**G**) Heat map of silhouette value measured with time-evolution of cellular processes effective contributions in segmentations based on time-average cellular processes (**E**), time-evolution of cellular processes (**F**), and large grid (**G**). (**H**) Histogram of silhouette value: blue for control segmentations, orange for region growing. Red vertical lines show silhouette value of label propagation results. Black vertical lines show silhouette value of regions smoothed by cellular Potts model. Dotted blue lines show threshold for the highest 0.05% of the control segmentations. Gray and cyan vertical line shows silhouette value of segmentation based on time-averaged cellular processes and large grid.

Also, the time-evolution of cellular processes was homogeneous inside the regions of the segmentation based on time-evolution of cellular processes, but not in segmentation based on time-average of cellular processes nor in the grid (Fig. 8E-G). The average silhouette value of segmentation based on time-evolution was higher than 99.995% of control segmentations (average silhouette for label propagation: 0.147, for smoothed regions: 0.1448, the maximum of the smallest 99.995% of controls average silhouettes: 0.124), while that of segmentation based on time-average was smaller than 5% of control segmentations (average silhouette for label propagation: 0.0125, for smoothed regions: -0.0385, the maximum of the smallest 95% of controls average silhouettes: 0.0241) (Fig. 8H).

Our tissue segmentation is designed to divide a tissue into regions homogeneous in a given property space, and the homogeneity of either tissue deformation rate or cellular processes in the segmentations based on each property demonstrated that the pipeline worked fine (Fig. 8B, F). However, it does not ensure the homogeneity of the regions in property spaces other than one based on which our segmentation was performed. Figure S4 shows heat maps of silhouette values measured in different property spaces. Even though the homogeneity in the regions differed among the different property spaces, the segmentations based on time-evolution of tissue deformation rate and cellular processes showed higher homogeneity than the others also in the property spaces of deformation rates due to cell divisions, cell rearrangements, and cell shape changes.

### 5.4 Cellular processes effective contributions inside the regions

We projected the regions divided based on the time-evolution of cellular processes onto the actual cell map, and found that the anterior and posterior regions corresponded to scutum and scutellum, and the middle boundary and lateral posterior regions corresponded to the scutum-scutellum boundary (Fig. 9). This result demonstrates that the obtained regions corresponded to the anatomical features, and cells were behaving differently between the anatomical regions. Note that the segmentations based on time-averaged tissue deformation rate or cellular processes did not match the anatomical features, indicating that the cells in the anatomical regions are actually regulated temporally.

**Fig. 9:**
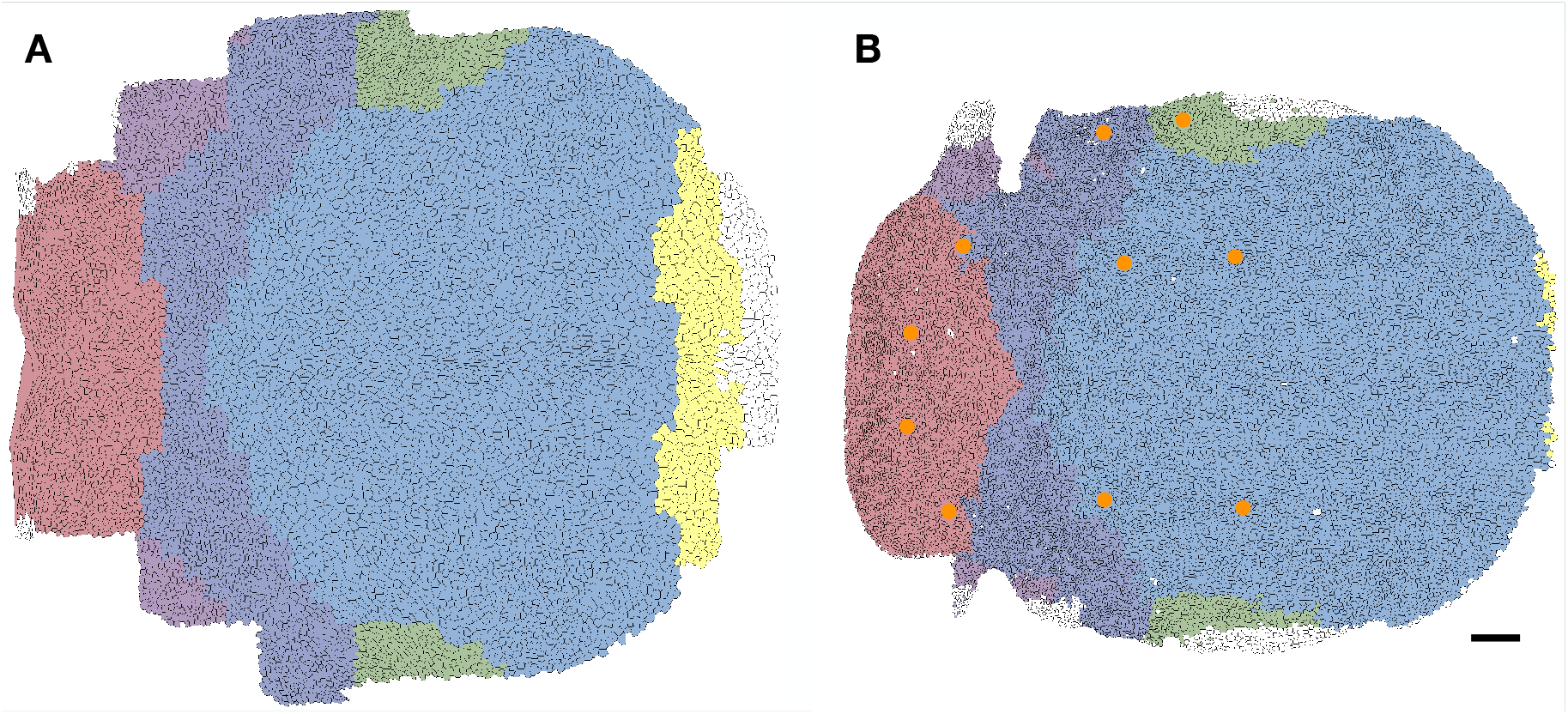
Projection of the segmentation onto the notum cells. The segmentations based on time evolution of cellular processes were projected. (**A**, **B**) The segmentation was projected onto the notum cells at 12 hr (**A**) and 32 hr APF (**B**), where regions were indicated by colors. The regions corresponded to scutum (pale blue and green), scutellum (red), scutum-scutellum boundary (dark blue and purple), and invaginated notum-neck boundary (yellow). The anatomical regions were identified according to positions of macrochaetae (orange circles in **B**). Scale bars indicate 50 *µ*m.

Figure 10 shows plots of cellular processes effective contributions average in each region of the segmentation based on time-evolution of cellular processes. The second peak of cell division was observed only in the posterior regions and the scutum-scutellum boundary region, consistent with the preceding studies with maps of number of cell divisions (Bosveld et al., 2012; Guirao et al., 2015), while we also found the first peak of cell division small in the lateral posterior region. Plots of average cellular processes differed from each other also among regions in the segmentation based on tissue deformation rate but less distinctive in the large grid segmentation (Fig. S5). Distances between the plots in Figure 10 were 0.65 *±* 0.16 and those for the segmentation based on tissue deformation rate were 0.63 *±* 0.20, larger than those for large grid segmentation (0.44 *±* 0.14). This result demonstrates that cellular processes in the obtained segmentations were more distinctive than those in the conventional grid.

**Fig. 10:**
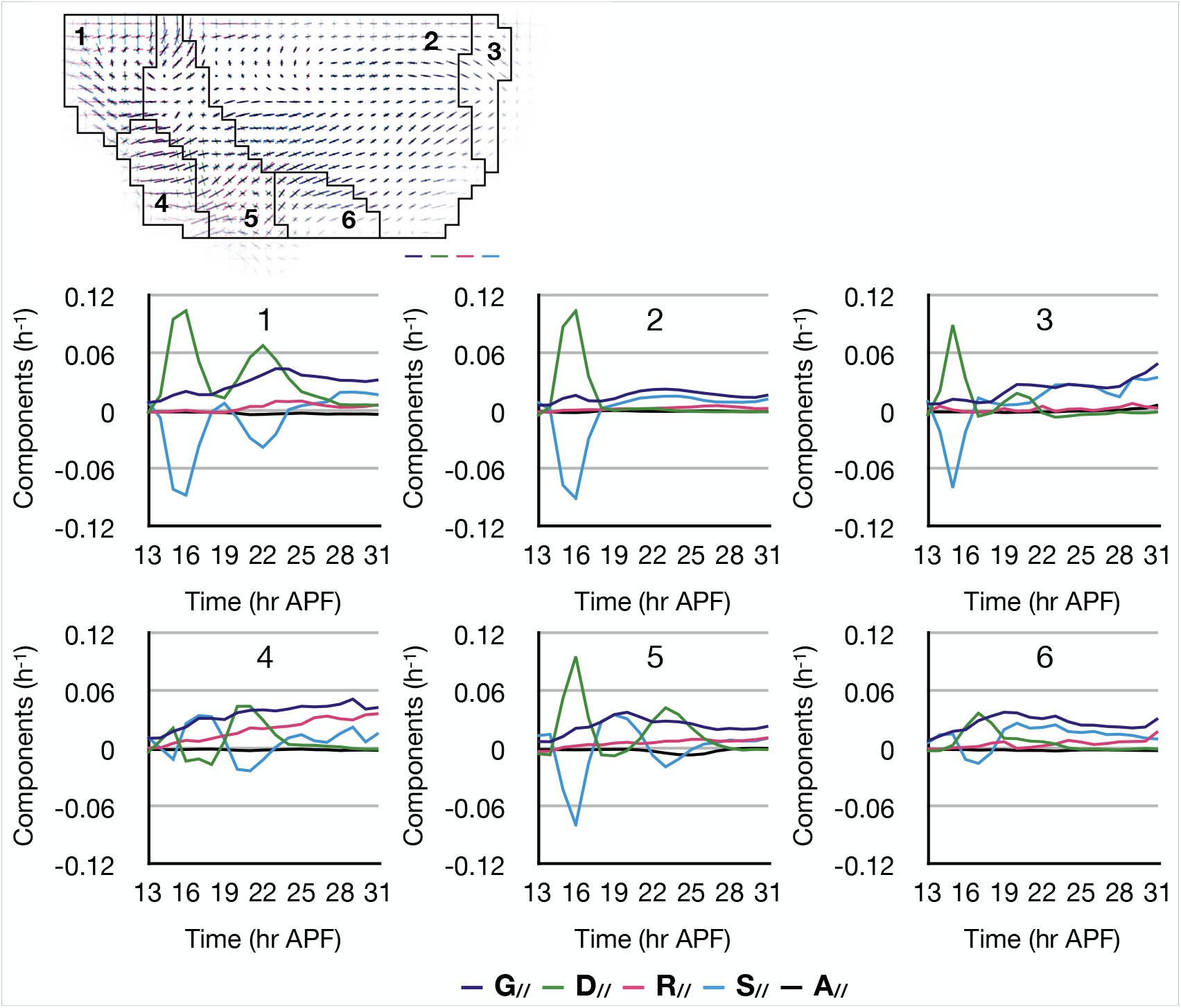
Cellular processes effective contributions inside the regions. (**A**) Tissue segmentation based on time-evolution of deformation rate, where two anterior subregions were merged. (**B**) Plots of cellular processes effective contributions averaged in each region of 1-5 in **A**.

### 5.5 Application to the morphogenesis of wing blade

To demonstrate the generality of our method to divide a tissue, we performed the same segmentation and analysis in the *Drosophila* pupa wing blade. During 15-32 hr APF, the wing blade is elongated in proximal-distal direction by a contracting wing hinge connected with the wing blade proximal side while its distal side is anchored to a cuticle via Dumpy (Etournay et al., 2015; Ray et al., 2015). The wing hinge contraction also narrows it in the anterior-posterior direction and induces shear strain in wing blade proximal anterior and posterior regions (Fig. 11A, B). We performed the tissue segmentation for the wing blade based on time-evolution of tissue deformation rate (Fig. 11C) and cellular processes (Fig. 11D) dividing into four regions. In both cases, the wing blade was divided into anterior, middle, posterior, and distal regions. All regions showed positive silhouette values (Fig. 11E, F), and their averages were significantly higher than the average silhouette values of control segmentations (Fig. 11G, H). Like the notum, dividing the wing blade into larger number of regions also subdivided already obtained regions (Fig. S3). Plots of the cellular processes effective contributions also showed distinctive patterns between the regions, where the cell division showed small contribution in the anterior region, the cell rearrangements dominated the tissue deformation around 26 hr APF in the anterior and posterior regions, and the cell shape changes showed two peaks around 17 and 22 hr APF in the distal region (Fig. 11I). Projection of the four regions onto the cells showed a difference between the regions and interveins, whereas the posterior region roughly corresponded to the proximal posterior intervein and the boundary between the anterior and distal regions corresponded L3 vein (Fig. S6).

**Fig. 11:**
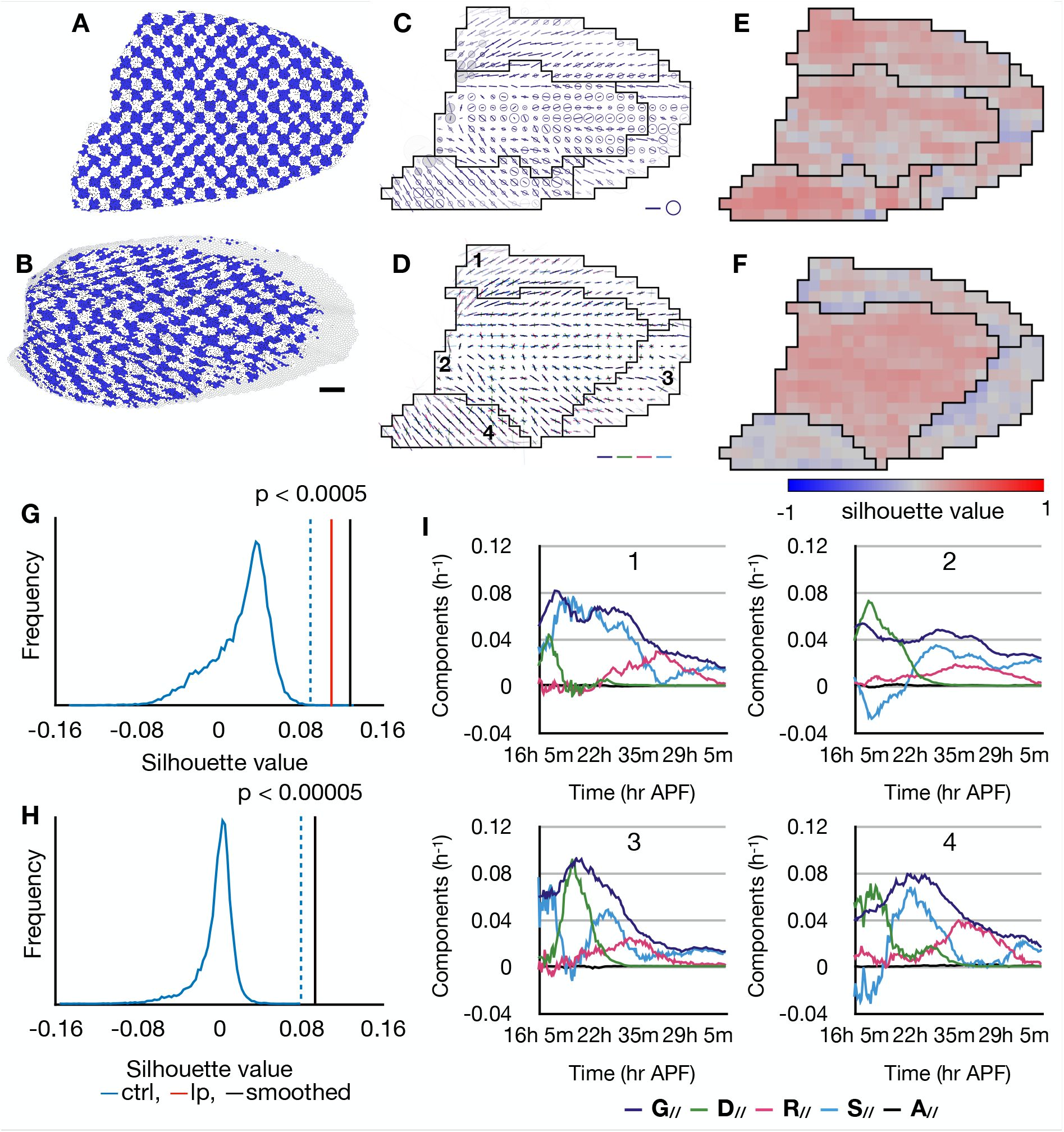
Segmentation of *Drosophila* wing blade into four regions. (**A**, **B**) Deformation of a wing blade from 15 hr (**A**) to 32 hr APF (**B**). Same representation ass Fig. 1 A, B. Cell patches are shown with blue and white check pattern. (**C**, **D**) Segmentation based on time-evolution of tissue deformation rate (**C**) and its heat map of silhouette value (**D**). (**E**, **F**) Segmentation based on time-evolution of cellular processes (**E**) and its heat map of silhouette value (**F**). (**G**, **H**) Histogram of average silhouette value of control segmentations for the property spaces of tissue deformation rate (**G**) and cellular processes (**H**). Red vertical lines show average silhouette values of label propagation results. Black vertical lines show average silhouette values of smoothed regions. Dotted blue lines show threshold for the highest 0.05% (**G**) and 0.005% (**H**) of the control segmentations. (**I**) Plots of cellular processes effective contributions averaged in each region of 1-4 in **D**.

## 6 Discussion

This study demonstrates that the pipeline of the region growing, the label propagation on the consensus matrix, and the boundary smoothing by cellular Potts model could divide a deforming heterogeneous tissue into homogeneous regions based on any prescribed quantity. Using this segmentation method, we divided the developing dorsal thorax and wing of *Drosophila* pupa based on their morphogenesis, and found regions with distinctive tissue deformation rate and underlying cellular processes.

The tissue segmentation based on morphogenesis differs from conventional image segmentation and cell segmentation algorithms in terms of quantity and supervision. Marking a human in a picture, an organ in histological image, or dividing cells in a microscopic image had been done manually, and thus results of the watershed algorithm or the artificial neural network could be supervised and corrected with the manual segmentations. On the other hand, the segmentation based on morphogenesis is hardly done manually for several reasons. First, the morphogenesis was quantified as multiple tensor fields with time evolution, and thus it is hard to visualize them in a 2D image for manual segmentation. Second, it is not easy to evaluate whether a given region actually corresponds to genetical/mechanical regulation of morphogenesis. Therefore we looked for a method which divides a tissue based on any prescribed quantity and returns regions with smooth boundaries. Region growing is a conventional and simple method of image segmentation, and requires a property space only to be metric. The varying results of the region growing were given to the label propagation and cellular Potts model to produce a single tissue segmentation, and the result was evaluated by region homogeneity boundary smoothness.

The notum segmentations based on time-evolution of tissue deformation rate and cellular processes effective contributions returned similar regions corresponding to the scutum, scutellum, and the boundary between them. By the tissue deformation rate, the scutum and scutellum regions were divided into medial and lateral subregions. Since the vector of effective contributions ignores the direction of deformation, the two subregions could be interpreted as regions of similar underlying cellular processes but deforming in different directions. On the other hand, the middle boundary region and the lateral region given by the cellular processes, both overlapped with the middle boundary region given by tissue deformation rate, could be interpreted as regions with different cellular processes but of similar tissue deformations. The wing blade was divided into anterior, middle, posterior, and distal regions based on both the tissue deformation rate and cellular processes, but the regions did not matched the wing veins pattern.

Silhouette analysis showed that the segmentations based on time-evolution of deformation rate and cellular processes included regions homogeneous in various property spaces, whereas the conventional grid segmentation included heterogeneous regions.

This method has some limitations. It cannot segment a small region; but such region might disappear during the boundary smoothing, resulting in disconnected regions in a final segmentation. Also, it is hard to determine the number of regions. We only know that a tissue might be over-segmented when it includes the small region. In a practical application, the tissue shall be segmented into various number of region with various minimum circularities, and one of them can be chosen by comparing it to gene expression patterns, analyzing cell behaviors inside each region, or other characteristics of interest.

In conclusion, we built a method to divide a tissue based on any prescribed property space. This allows an application to a study of spatial regulation of various processes, where the property space should be chosen for the process of interest. For example, to study a spatial regulation of cell division orientation, the property space may be prepared from, instead of the local deformation rate and cellular processes, the tensor field of cell division and known regulating factors such as cell shape, localization of planar cell polarity proteins, and tension on cell-cell interface, and then resultant regions can be compared with genes expression patterns. Also, this method is not dependent on how the morphogenesis was quantified, and one can include rotational movement by anti-symmetric strain rate tensors, or 3D deformation by using voxels instead of pixels.

## 7 Methods

### 7.1 Quantification tools

This section describes in details the following quantification tools. Morphogenesis data result from the quantification of local tissue deformation rate and underlying cellular processes as described in (Guirao et al., 2015). The similarity between two tensors is quantified by the standard Euclidean metric. The homogeneity of a quantity within a given region, i.e. the similarity between measurements of this quantity within a region, is measured by silhouette, a standard tool of cluster analysis. For a measurement of similarity between tissue segmentations, we use the Rand index, which indicates how well two data clusterings agree.

#### 7.1.1 Quantification of tissue deformation and cellular processes

Quantification of local tissue deformation and underlying cellular processes was performed in (Guirao et al., 2015). Briefly, *Drosophila* nota expressing GFP-tagged E-cadherin were imaged. The notum movies were split in a grid (with patches about 20 *µ*m width) at the first frame (Fig. 1C), 12 hr after pupa formation (APF). The local deformation rate and the cellular processes were measured in each cell patch through the development, as follows.

Epithelial cell contours were detected automatically using watershed algorithm, cells were tracked, adjacencies between cells were listed, and relative positions of adjacent cell centers were recorded. The tissue deformation rate, denoted by the symmetric tensor **G**, was obtained from changes of relative positions between neighbor cells over 20 hr from 12 hr APF to 32 hr APF, or over 2 hr at each time point when recording the time evolution. The tissue deformation rate **G** was then decomposed into cell shape change **S** and deformation accompanied by change of cell adjacency, which was further divided into cell division **D**, cell rearrangement **R**, and cell delamination **A**, which are symmetric tensors too.

In a collection of cells where the total deformation is driven completely by the four fundamental cellular processes, the tensors are in a balance equation,

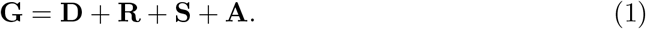

The scalar product of two tensors **Q** and **Q***^!^* in dimension *d* is defined as:

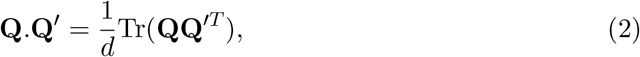

and the unitary tensor **u_G_** that is aligned with **G** is given by

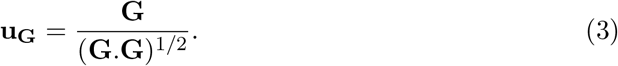

Since the scalar product (Eqn. 2) is a bilinear operation, multiplying **u_G_** by a tensor, the operation .**u_G_** : **Q** *→* **Q***_//_*, retains the balance between the tissue deformation rate and the cellular processes in Equation 1 while converting them to scalar magnitudes:

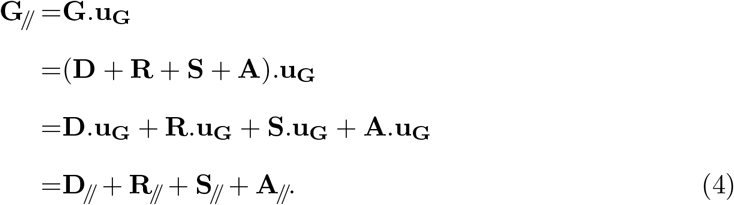

The scalar **G***_//_* represents the local magnitude of tissue morphogenesis, and **D***_//_*, **R***_//_*, **S***_//_*, and **A***_//_* represent the effective contributions of the cellular processes to the tissue morphogenesis. When a cellular process produces an anisotropic deformation in the same direction as that of tissue, e.g. cells divided in the same direction as tissue elongation, the scalar product between them returns a positive value, while it returns a negative value when a cellular process counteracts tissue deformation.

#### 7.1.2 Metric

Similarity of morphogenesis between different cell patches was defined as follows.

For expansion/contraction of area (isotropic deformation), similarity was given by difference in expansion/contraction rates.

Similarity of anisotropic deformation was given by a distance between two tensors **Q** and **Q’**,

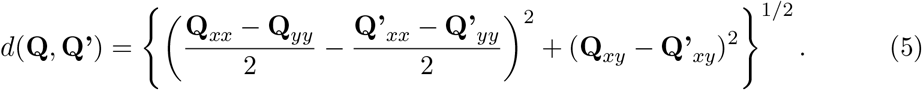

For tensors with time-evolution **Q**(*t*) and **Q’**(*t*), distance was given by a sum of the distance at each time point,

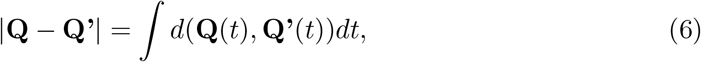

as an analogy to distance between functions.

For the composition of cellular processes, the tensors of cellular processes were converted to effective contributions and combined into a vector (**G***_//_,* **D***_//_*, **R***_//_*, **S***_//_*, **A***_//_*). A distance between two vectors was given by Euclidean distance, the square root of the sum of the square of the differences between corresponding elements, and a distance between vectors with time-evolution *v*(*t*) and *v^!^*(*t*) was given by a sum of the distance at each time point,

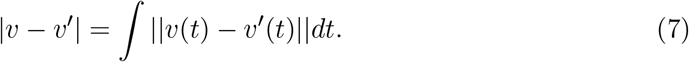

#### 7.1.3 Silhouette and bootstrap

Silhouette quantifies clustering results, indicating how well an object resembles other objects inside its own cluster (Rousseeuw, 1987). Assume that *n* objects *{p*_1_*, p*_2_*, . . . , p_n_}* are partitioned into *k* clusters *{C*_1_*, C*_2_*, . . . , C_k_}*. For an object *p_i_ ∈ C_I_* , we can compute the average distance *a*(*p_i_*) from *p_i_* to all other objects in *C_I_* . For *J* ≠ *I*, we can also compute the average distance *d*(*p_i_, C_J_* ) from *p_i_* to all objects in *C_J_* , and select the smallest of those, denoted by *b*(*p_i_*) = min*_J_*_=_*_I_ d*(*p_i_, C_J_* ). The silhouette value *s*(*p_i_*) is obtained by combining *a*(*p_i_*) and *b*(*p_i_*) as follow:

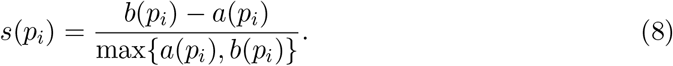

By this definition, *−*1 *≤ s*(*p*) *≤* 1, where *s*(*p*) large and close to 1 indicates that *p* is similar to other objects in the same cluster, while negative *s*(*p*) indicates that there is another cluster whose objects are more similar to *p* than objects in the cluster containing *p*.

We took the average silhouette value over all points (cell patches) as a measurement of homogeneity of a given segmentation. For significance test, tissue was segmented randomly (see below) 20,000 times into a given number, and we got thresholds above which the highest 5%, 0.5%, or 0.005% of the average silhouettes were found. The average silhouette of given regions were compared to those of the control segmentations with the same number of regions.

#### 7.1.4 Adjusted Rand index

For a measurement of similarity between tissue segmentations, we use the Rand index, which indicates how well two data clusterings agree ; its value is 0 if the clusterings entirely disagree and 1 if they entirely agree. Its corrected-for-chance version is a more meaningful quantity, called the adjusted Rand index (ARI): it is the Rand index compared with its value expected for the random case, and its value can be negative.

We compute the adjusted Rand index with the permutation model (Hubert and Arabie, 1985). Given two clusterings *A* = *{A*_1_*, . . . , A_k_}* and *B* = *{B*_1_*, . . . , B_m_}* of *N* elements, the contingency table *τ* = (*n_ij_* )*_k×m_* is made where *n_ij_* = *|A_i_ ∩ B_j_|*. The Rand index between *A* and *B*, RI(*A, B*) is

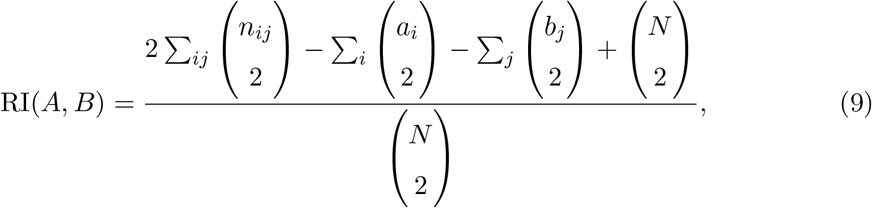

where *a_i_* = Σ*_j_ n_ij_* and *b_j_* = Σ*_i_ n_ij_* , and for the random case the expected Rand index 𝔼[RI(*A, B*)] is

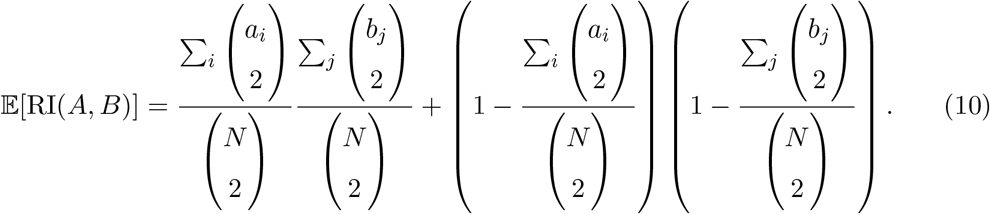

Finally, the adjusted Rand index ARI(*A, B*) is

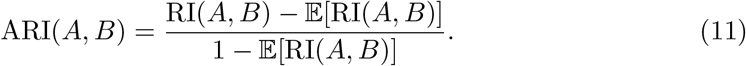

### 7.2 Tissue segmentation pipeline

The pipeline was implemented by custom Matlab scripts, in three steps (Fig. 1C). The Matlab scripts are available at GitHub (https://doi.org/10.5281/zenodo.4270726). Also, we provides pseudo codes of them in the supplementary materials.

#### 7.2.1 Region growing tissue segmentation

The initial tissue segmentation first defines a metric of similarity between cells, and then a tissue is divided into regions containing similar cells. This approach was inspired by a study segmenting mouse heart based on cell polarity (Le Garrec et al., 2013). On the assumption that expression patterns of genes responsible for morphogenesis make connected regions, and to study physical interactions between the regions, we aimed at getting connected regions.

The algorithm *Region growing* (Adams and Bischof, 1994; Ma et al., 2010) is an image segmentation method using a process similar to *k*-means clustering, starting from randomly given seeds (corresponding to “means” in *k*-means clustering), segmenting an image with the seeds followed by update of the seeds within the regions, and iterating this process until convergence (Fig. 2A). The tissue segmentation is done by growing regions from the seeds collecting pixels adjacent to the growing regions, and so the resultant regions are connected.

Initial seeds were randomly chosen from data, and regions were expanded by adding a pixel (cell patch) adjacent to a region and the most similar to the seed of the region in the property space one by one until all pixels were assigned to one of the regions. The seeds were updated to pixels the closest to centroids of the regions, averages of the regions in the property space were given as property of the seeds, and then regions were expanded again from the seeds. These region expansions and seed updates were iterated until convergence was reached.

#### 7.2.2 Preparation of control segmentations

The control segmentations were made using an algorithm similar to the region growing but ignoring the similarity between points. From randomly given seeds, regions were expanded by adding a pixel adjacent to a region, where the added pixel was chosen randomly from all adjacent pixels, until all pixels were assigned to one of the regions. Therefore any obtained region is connected.

#### 7.2.3 Label propagation on a consensus matrix

To merge multiple segmentation results into a single one independent on the metric, we use label propagation algorithm on a consensus matrix, which takes multiple partitions and returns a consensus partition which is the most similar to all partitions (Lancichinetti and Fortunato, 2012; Raghavan et al., 2007).

For a division of *n* points, independent 50 trials of region growing were converted to a consensus matrix, whose entry at *i*-th row and *j*-th column indicates a frequency of partitions in which *i*-th point and *j*-th point were in the same cluster. The entries lower than a given threshold were set to 0. The label propagation started by assigning to each point a different label. Then the label of randomly chosen *i*-th point was updated to one that was the most weighted by the consensus matrix, where *ij* element gave the weight to a label of *j*-th point. The label update was iterated until convergence. The threshold for the consensus matrix was scanned between 20-80% so that a resultant partition contained the same number of regions as the initial partitions.

#### 7.2.4 Cellular Potts model for boundary smoothing

To smooth the consensus region boundaries while preserving region area and homogeneity, we use the cellular Potts model, in which a cellular structure is numerically simulated in a square lattice, where each cell is a set of pixels. The system energy depends on cell shapes, and the pattern is updated in an iteration to decrease the energy, with some fluctuation allowance (Graner and Glazier, 1992). In the simplest and common two-dimensional form, the energy *H* arises from total perimeter length *P* (with line energy *J* ) and constraint on each region area *A* (with compressibility *λ*); decreasing it results in smoother regions with preserved area *A*_0_, removing small protrusions and disconnected regions. In this study, we also included the silhouette *s* to account for the region homogeneity, with a weight coefficient *h*:

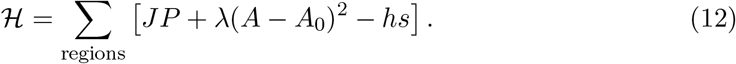

The coefficients *J* , *λ*, and *h* were screened as described below.

When updating the label for a randomly selected pixel *a*, a target label was randomly selected from neighbors of *a*, and then Hamiltonian change was calculated. Label of *a* was updated to the target label with probability *min*(1*, e^−^*^Δ^*^H/T^* ), where Δ*H* denotes change of *H* by the change of label of *a*, and *T* is the fluctuation allowance. In the present application, the updates of labels were iterated 50 times. For resultant regions, a circularity *C* was calculated, where it was defined as *C* = 4*π×*area*/*perimeter^2^ (Bosveld et al., 2016). The parameters *J* , *λ*, *h*, and *T* were screened for resultant regions with the highest homogeneity and circularity larger than a given threshold value. The screening was performed in a pairwise testing manner on a grid, and grid was converged to the highest homogeneity. With the screened parameters, the boundary smoothing was iterated for 50 times, and the results were integrated again by the label propagation on a consensus matrix algorithm.

#### 7.2.5 Cellular Potts model for tissue compression simulation

For the simulation of cells in a compressed tissue, the cellular Potts model was implemented by custom Matlab scripts which are available at GitHub (http://doi.org/10.5281/zenodo.5016684).

It is simulated on a torus surface so the cells made a periodic pattern. The initial configuration was prepared by the Voronoi tessellation from randomly scattered 600 points. The points were randomly selected from 864 *×* 150 pt^2^ plane in a sequential manner so that all points were away from the others at least 10 pt. Cells on the left side were assigned the low line energy *J*_soft_ = 1 and cells n the right side were assigned the high line energy *J*_stiff_ = 4. All cells were assigned same compressibility *λ* = 1. For the compression, the initial configuration was transformed into a 480 *×* 270 image. Then a dynamics of the cells was simulated for 975,000 updates.

To be processed by the tools developed in (Guirao et al., 2015), images of the cells were projected on a plane and converted to cell segmentation. Since all pixels belonged to one of the cells in the cellular Potts model and the cells were not separated by boundary (watershed) pixels, we first labeled the pixels with *perimeter* or *inner*, where the *perimeter* pixels were adjacent to different cells and the *inner* pixels were enclosed by the *perimeter* pixels. Then the *perimeter* pixels were re-labeled one-by-one to *inner* so that all *inner* pixels were connected in each cell, all *perimeter* pixels were adjacent to *inner* pixels of the same cell, and no *inner* pixel was adjacent to *inner* pixel in different cells. The labels were updated as much as possible, and remained *perimeter* pixels were taken as the boundary.

## Acknowledgement

The authors thank Floris Bosveld for imaging, Stéphane U. Rigaud for cell segmentation and tracking, Yohanns Belläıche for useful advices, and all team members for insightful discussions.

## 9 Author contributions

Quantification of morphogenesis: B.G.; Methodology: S.Y., F.G.; Programming and implementation: S.Y.; Writing and editing: S.Y., F.G.

---

**Fig. S1:**
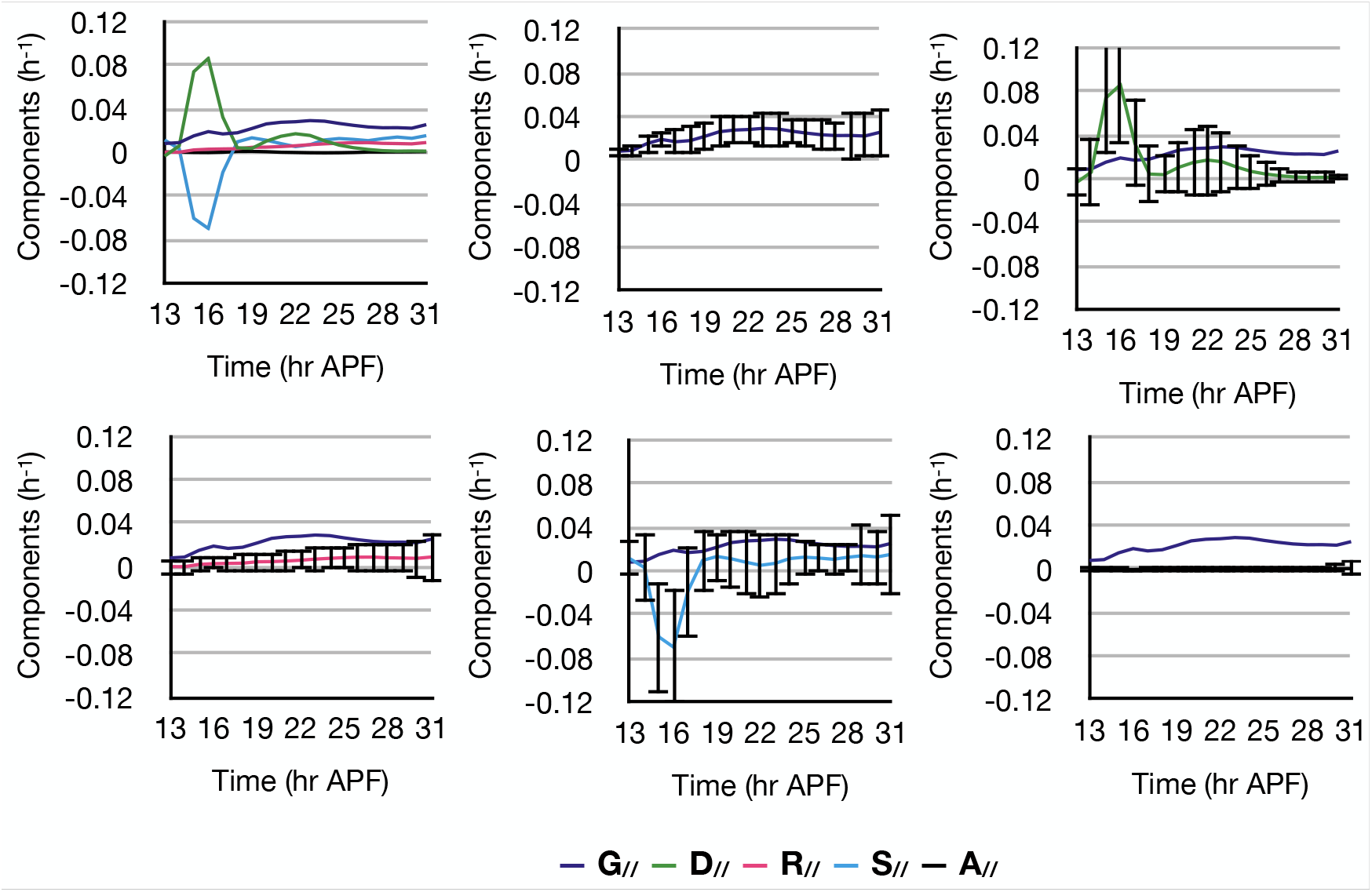
Variance of effective contribution of cellular processes at each time point. Plots show time evolution of effective contribution of cellular processes in the *Drosophila* notum and standard deviation of them.

**Fig. S2:**
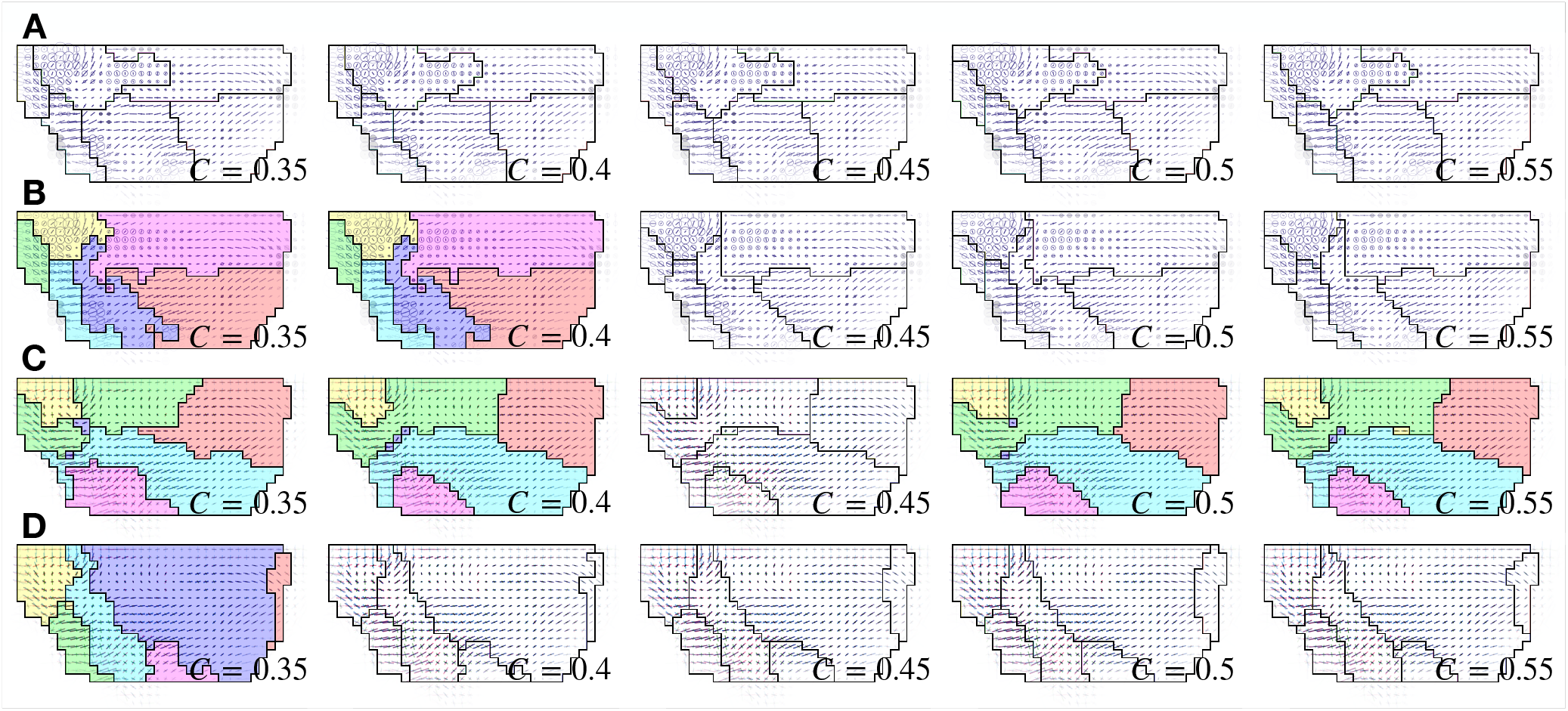
Boundary smoothing with various minimum circularities. The *Drosophila* notum was divided based on time-average tissue deformation rate (**A**), time-evolution of tissue deformation rate (**B**), time-average cellular processes effective contributions (**C**), and time-evolution of cellular processes effective contributions. They were smoothed with the minimum circularity *C* ranging from 0.35 to 0.55. Some of them were colored for visibility.

**Fig. S3:**
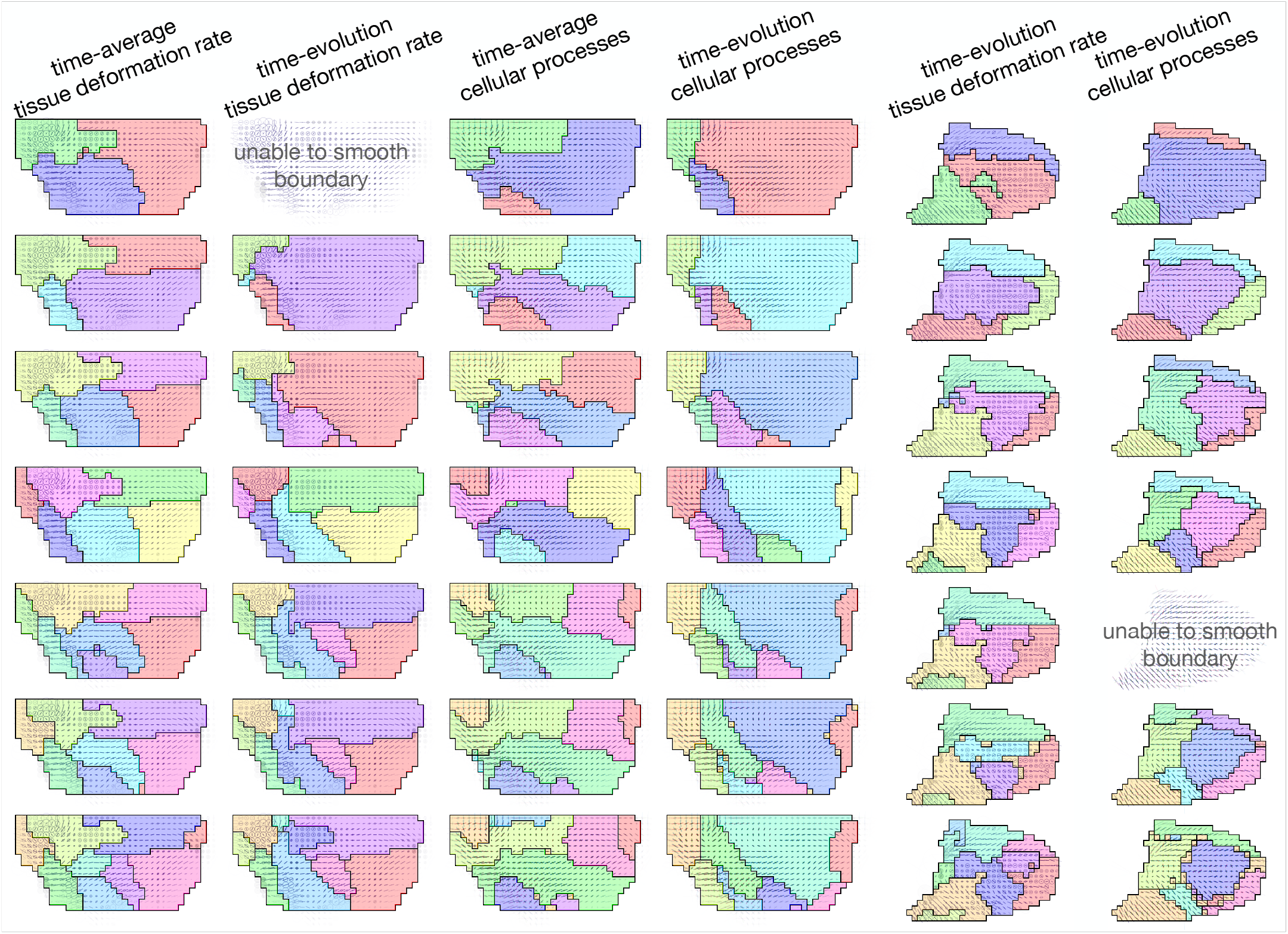
Segmentations in different number of regions. Dorsal thorax (first to fourth columns) and wing blade (fifth and sixth columns) were divided into 3-9 regions. First column: segmentations based on time-average tissue deformation rate. Second column: segmentations based on time-evolution of tissue deformation rate. Third column: segmentations based on time-average cellular processes effective contributions. Fourth column: segmentations based on time-evolution of cellular processes effective contributions. Fifth column: segmentations based on time-evolution of tissue deformation rate. Sixth column: segmentations based on time-evolution of cellular processes effective contributions. The tissues were divided into 3 to 9 regions (from top to bottom rows). The regions were colored for visibility. When the number was too large and a result of the initial label propagation included a too small region, the small region tended to disappear in the cellular Potts model smoothing, and thus the final label propagation tried to integrate regions fewer than the final segmentation, sometimes resulted in undesired disconnected regions (third column bottom row and fourth column sixth row). For dividing the dorsal thorax into three regions based on time-evolution of tissue deformation rate and wing blade into seven regions based on time-evolution of cellular processes effective contributions, it failed to screen the parameters (the screening algorithm pursued to a too low temperature which would freeze any change in the cellular Potts model, second column first row and sixth column fifth row).

**Fig. S4:**
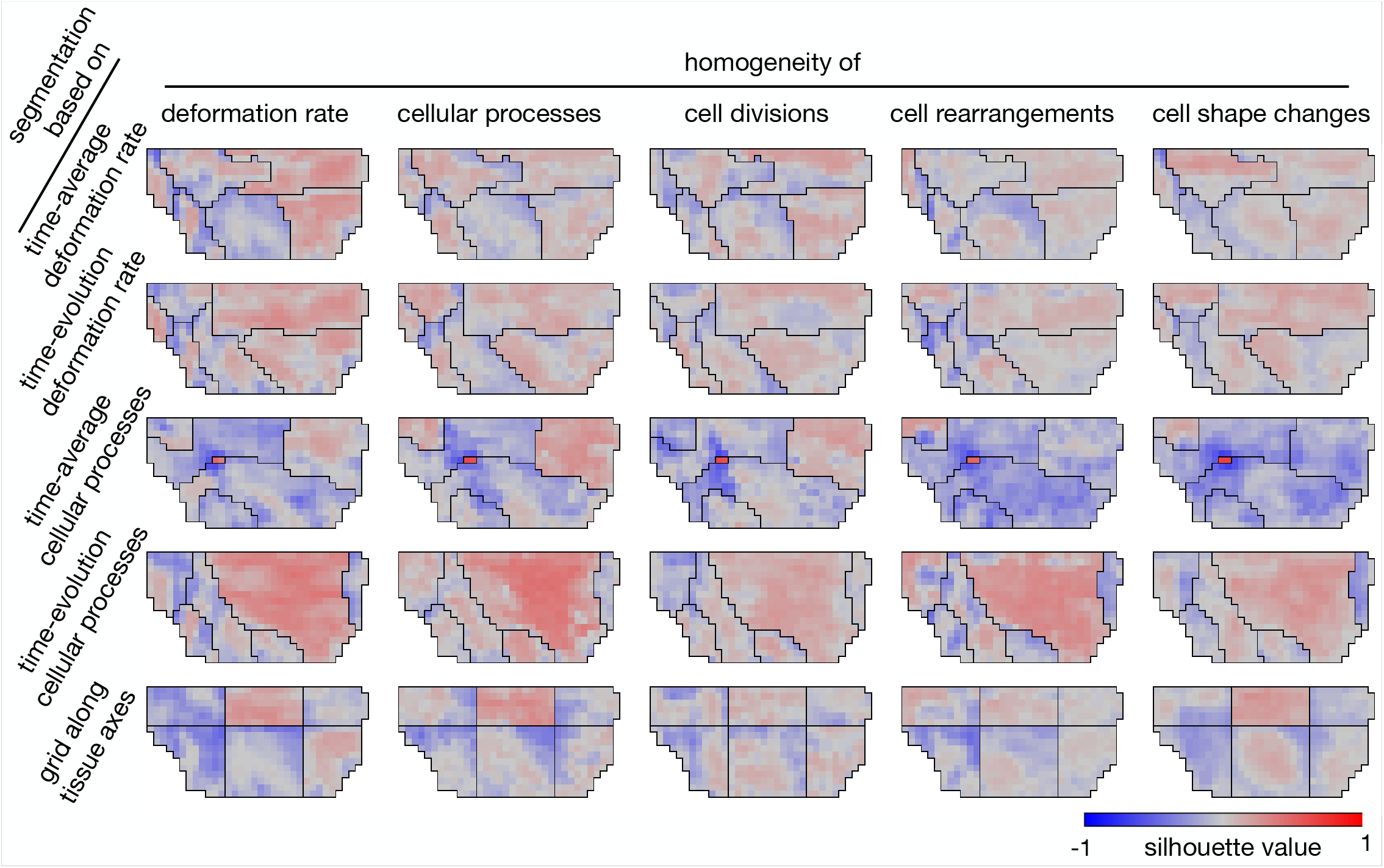
Heat maps of silhouette value. First row: segmentation based on time-average tissue deformation rate. Second row: segmentation based on time-evolution of tissue deformation rate. Third row: segmentation based on time-average cellular processes effective contributions. Fourth row: segmentation based on time-evolution of cellular processes. Fifth row: conventional segmentation of large grid parallel to tissue axes. First column: silhouette values measured in the property space of time-evolution of deformation rate. Second column: silhouette values measured by time-evolution of cellular processes. Third column: silhouette values measured by time-evolution of cell divisions. Fourth column: silhouette values measured by time-evolution of cell rearrangements. Fifth column: silhouette values measured by time-evolution of cell shape changes.

**Fig. S5:**
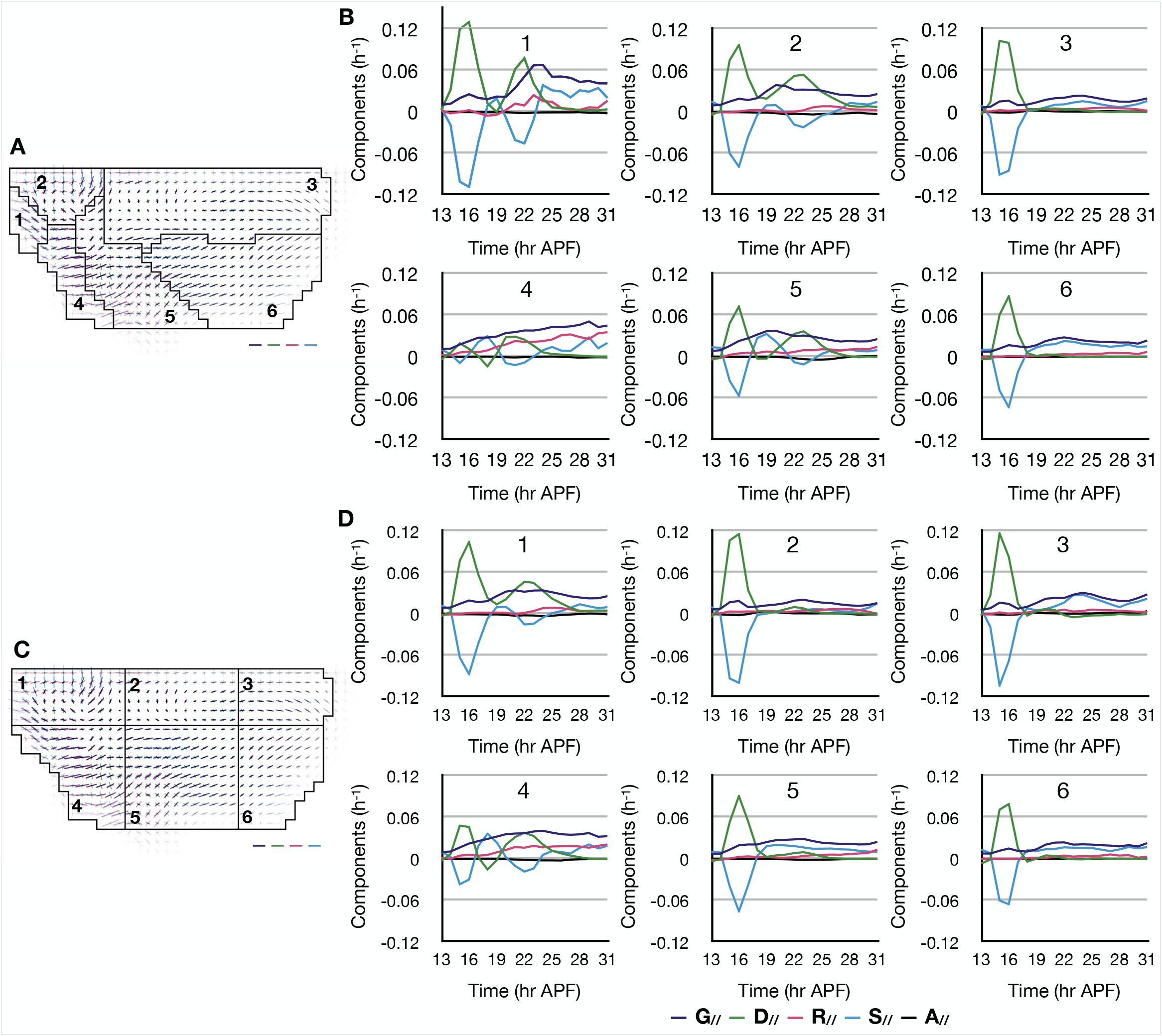
Plots of cellular processes in the segmentations based on time evolution of tissue deformation rate and the conventional large grid. (**A**, **B**) The tissue segmentation based on time-evolution of tissue deformation rate (**A**) and plots of cellular processes effective contributions averaged in each region (**B**). The numbers indicate the regions. (**C**, **D**) The large grid (**C**) and plots of cellular processes in each region (**D**). Scale bars in **A** and **C** indicate deformation rate 0.02 h*^−^*^1^ with colors for tissue and cellular processes.

**Fig. S6:**
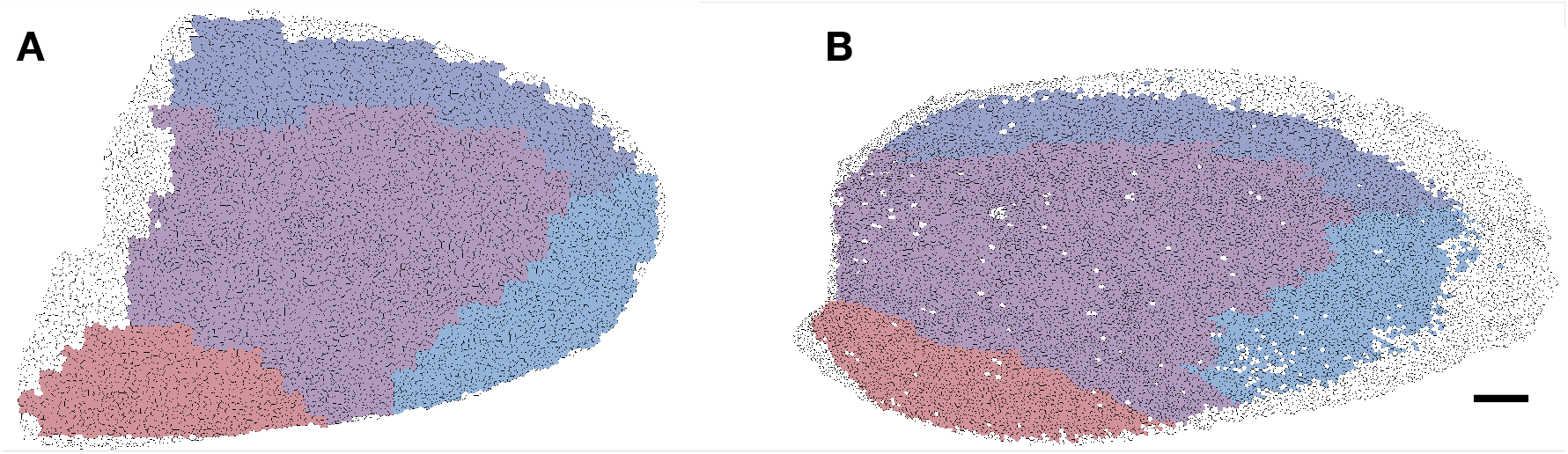
Projection of the segmentation onto the wing blade cells. The segmentations based on time evolution of cellular processes were projected. (**A**, **B**) The segmentation was projected onto the wing blade cells at 15 hr (**A**) and 32 hr APF (**B**), where the regions were indicated by colors. Scale bars indicate 50 *µ*m.

**Movie 1:** Cell rearrangements and cell shape changes after tissue compression. The cells with low and high surface tension were colored gray and yellow respectively. The cellular Potts model was run on an image of 480 *×* 270 lattice and included 600 cells. The movie is 7 fps and there were 5000 updates between the frames.

## 1 Pseudo codes for tissue segmentation algorithms

In below pseudo codes show algorithms of the automatic tissue segmentation. Matlab custom functions and framework developed for this study are available at GitHub (http://doi.org/10.5281/zenodo.3626111 and http://doi.org/10.5281/zenodo.5016684). For details of the functions and framework, see its README file and comments in the codes.

### 1.1 Region growing algorithm

Algorithm 1 shows a pseudo code of the region growing image segmentation in Matlab-like syntax. It divides a bitmap image stored in a data object *dataMap*. In the algorithm, a number of regions, a limit to update the seeds, and a metric are given as parameters. With the parameters, supporting objects *seedList*, *meanList*, *regionsList*, *meter*, and *seeder* are allocated and initialized. The seedList, meanList, and regionsList are instances of data object with a property *var* representing seeds and means of regions and regions, shared among the supporting objects. The meter is an object measuring distance between the mean of region and a point adjacent to the region. A method *measure* returns the distance measured by the given metric. The seeder is an object choosing seeds of regions. Methods *initalSeeds* and *initialMeans* return indices of randomly chosen points and their values. Once the dataMap was divided into regions, methods *newSeeds* and *newMeans* return indices of points at center of the regions and mean values of the regions. A method *initalQueue* returns an array where its element represents a point adjacent to one of the seeds and holds the region and distance to the region’s mean value. Inside a loop, a point in the queue with the smallest distance to the region’s mean value is added to the region, and points adjacent to the point, returned by a method *neighborsOfPoint* of dataMap, are added to the queue.

In our tissue segmentation, a Matlab custom function *run_region_growing* () iterates this algorithm for given time, returning a stack of resultant partitions.

### 1.2 Label propagation on a consensus matrix

Algorithm 2 shows a pseudo code of the label propagation. It divides *N* objects into clusters based on an *N × N* consensus matrix *M* whose rows and columns correspond to the objects, and an element *m_ij_* represents the frequency at which the *i*-th and *j*-th objects were included in a cluster among given clustering results. A parameter *t_M_* indicates a threshold value, where elements in *M* smaller than *t_M_* are ignored in the label propagation.

In the tissue segmentation, 50 results of region growing were converted to the consensus matrix and given to a Matlab custom function *run_label_propagation*() implementing the label propagation. The number of resultant regions is influenced by *t_M_* , and thus a Matlab custom function *run_cm_thresholding_lp*() screens *t_M_* values so that *run_label_propagation*() returns the same number of regions with the given partitions.

#### Algorithm 1: Region growing algorithm

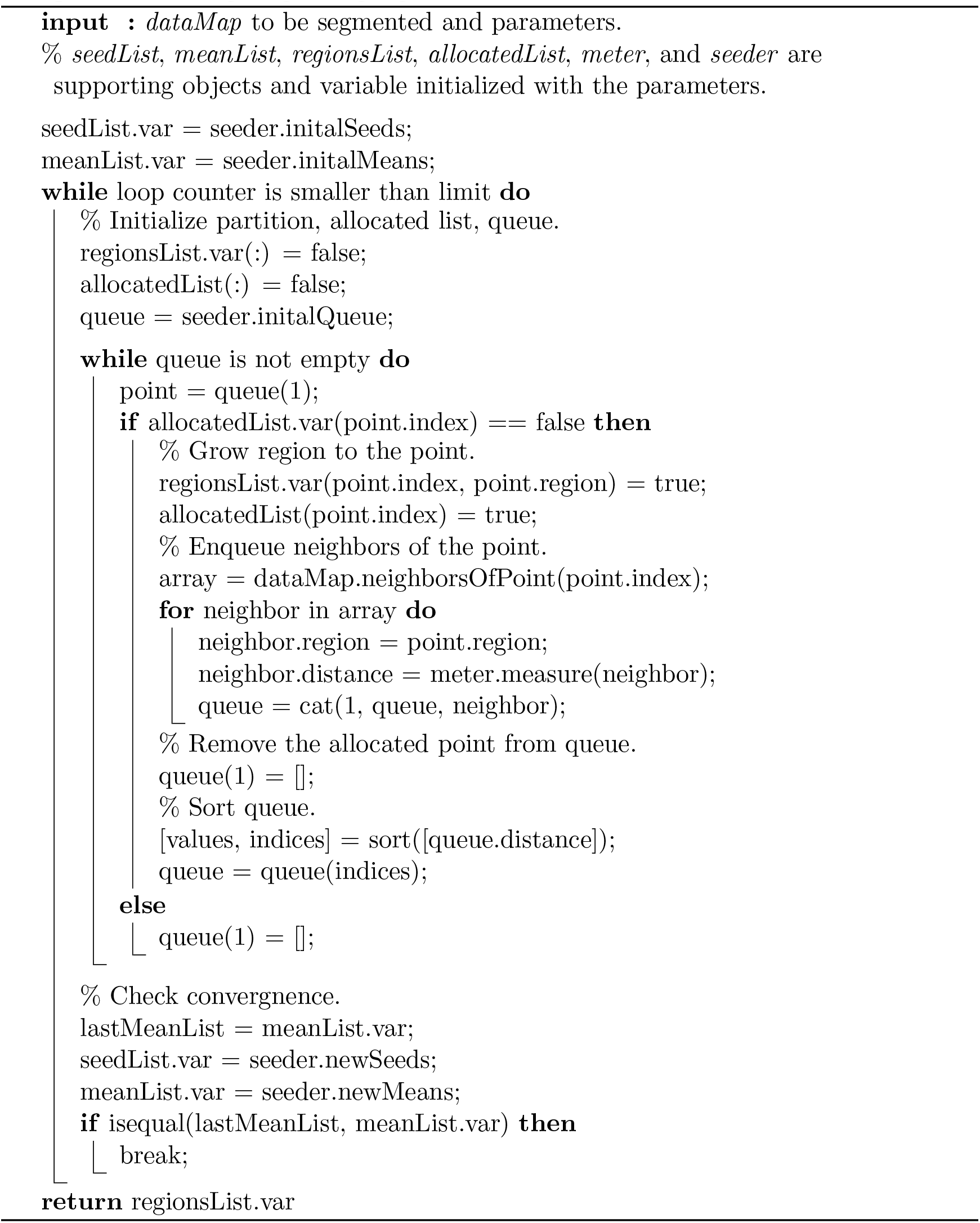

#### Algorithm 2: Label propagation

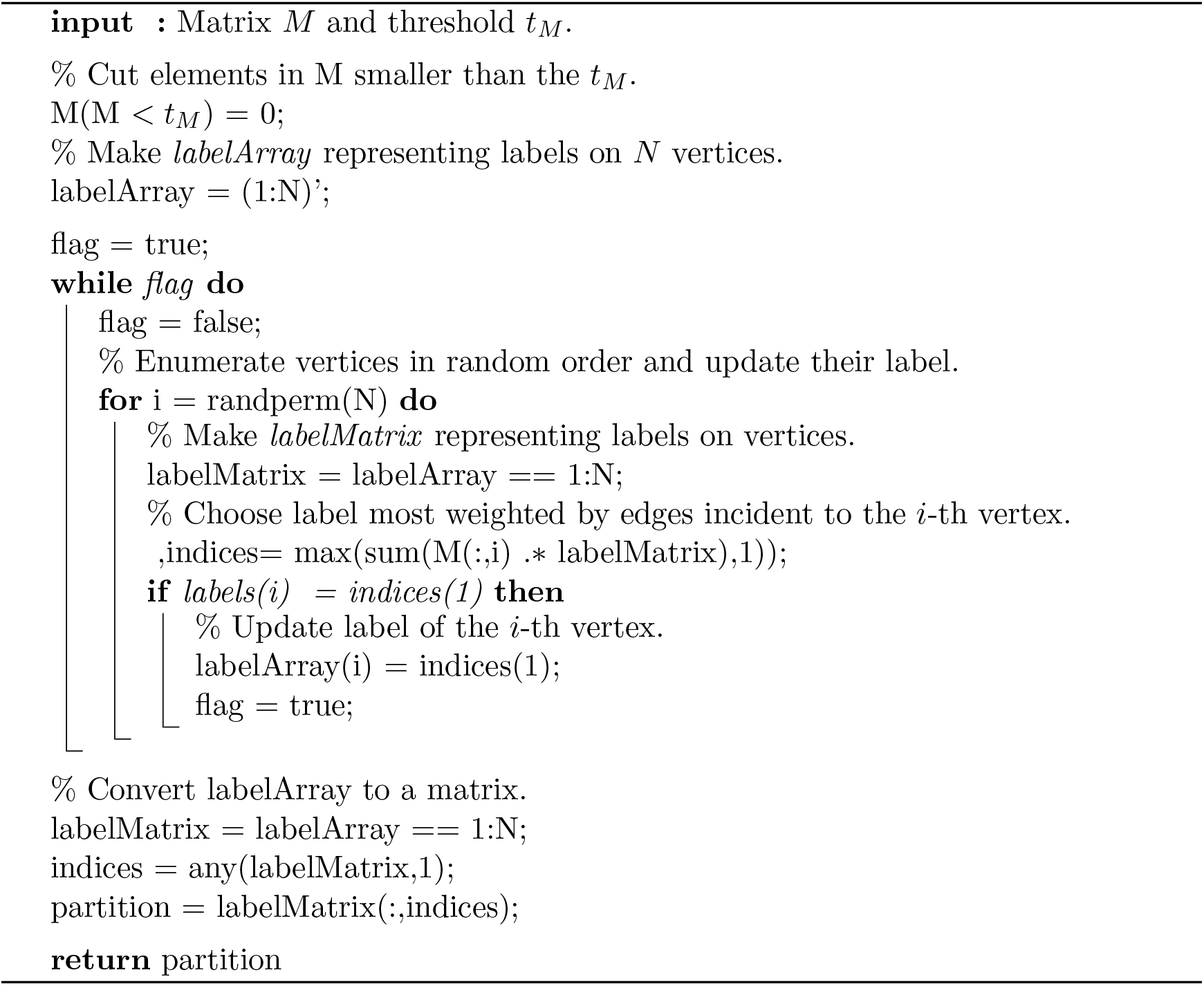

### 1.3 Cellular Potts model

Algorithm 3 shows a pseudo code of the cellular Potts model. It simulates a deformation of regions (partition of dataMap) by giving small fluctuations. In the algorithm, an array of function handles, coefficients to combine the functions results, the system temperature, and the number of label updates are given as parameters. With the regions and parameters, supporting objects *analyser* and *dict* are allocated and initialized. The functions in the array calculate system energy with *analyser* and *dict*. For each fluctuation, one of points at regions rim returned by analyser *rim points* is selected randomly, and a label of neighboring points is also selected randomly and copied. Connectedness of a region is checked locally, with a coordinate of neighboring points returned by dataMap *coordinates*.

In the tissue segmentation, a Matlab custom function *run_CPM_smoothing* () implement this algorithm with energy functions combining area constraint, surface tension, and total silhouette value. The coefficients and temperature influence resultant regions, and thus a Matlab custom function *run_CPM_fitting* () screens the parameters so that *run_CPM_smoothin*() returns smoothed regions with a circularity larger than the given value and the total silhouette value as large as possible.

#### Algorithm 3 Dummy node preparation phase

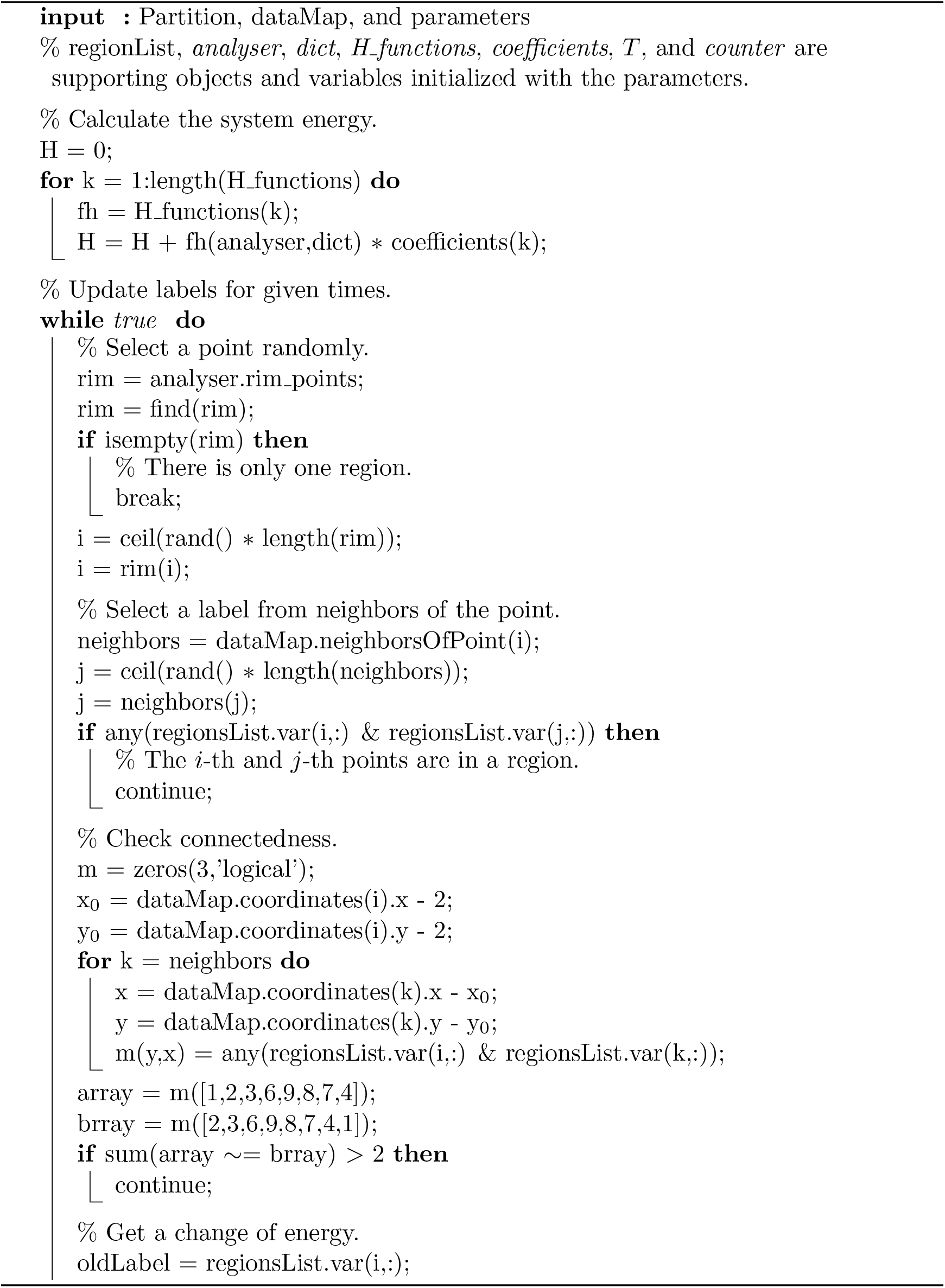
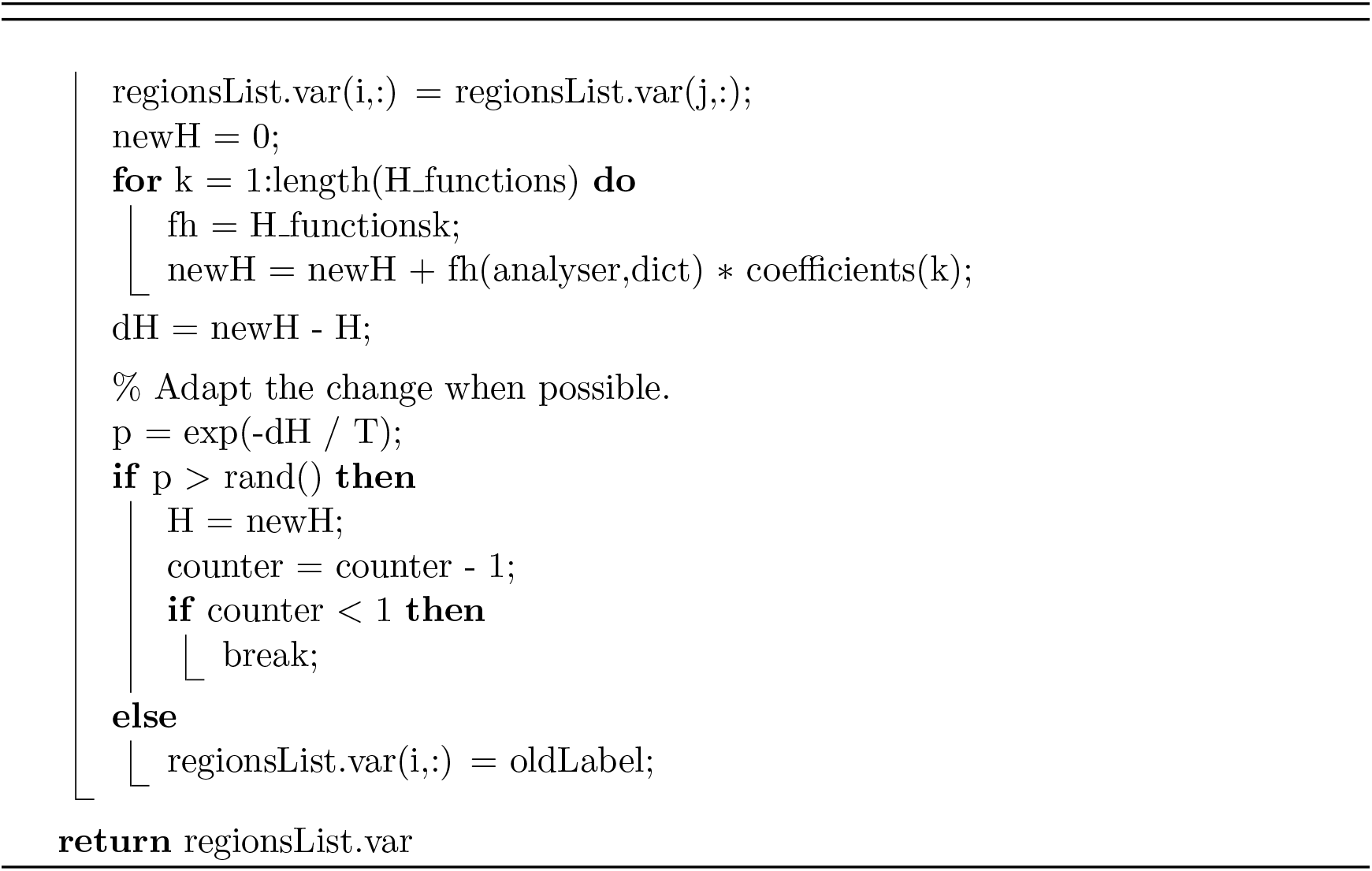

